# Scoring gene importance by interpreting single-cell foundation models

**DOI:** 10.1101/2025.06.14.659567

**Authors:** Maxwell P. Gold, Miguel Reyes, Nathaniel Diamant, Tony Kuo, Ehsan Hajiramezanali, Jane W. Newburger, Mary Beth F. Son, Pui Y. Lee, Gabriele Scalia, Aicha BenTaieb, Sharookh B. Kapadia, Anupriya Tripathi, Héctor Corrada Bravo, Graham Heimberg, Tommaso Biancalani

**Affiliations:** gRED Computational Sciences, Genentech, South San Francisco, California; gRED Research Biology, Genentech, South San Francisco, California; Roche Informatics, F. Hoffmann-La Roche, Mississauga, Ontario, Canada; Department of Cardiology, Boston Children’s Hospital, Boston, Massachusetts; Department of Pediatrics, Harvard Medical School, Boston, Massachusetts; Division of Immunology, Boston Children’s Hospital, Harvard Medical School, Boston, Massachusetts

## Abstract

Determining a gene’s functional significance within a cellular context has long been a challenge, as absolute expression level is an unreliable indicator. We introduce SIGnature, a framework for scoring gene importance by leveraging attributions derived from single-cell RNA-sequencing (scRNA-seq) foundation models. Attribution scores reduce technical noise, emphasize regulatory genes, and facilitate cross-dataset comparison – a core challenge for scRNA-seq analyses. We developed the SIGnature package as a tool for generating and querying attributions, enabling rapid gene set searches across massive scRNA-seq atlases. We demonstrated its utility using the MS1 monocyte signature, a poorly understood gene program activated in severe COVID-19 and sepsis. Searching 400 studies revealed novel associations between the MS1 signature and multiple hyperinflammatory conditions, including Kawasaki disease. Experimental validation confirmed Kawasaki disease patient serum induces the MS1 phenotype. These findings highlight that SIGnature can uncover shared mechanisms across conditions, demonstrating its power for large-scale signature scoring and cross-disease analysis.

## Main

A gene’s absolute expression level is not a reliable indicator of its functional significance within a single cell. For instance, key regulators like transcription factors are typically lowly expressed, while highly abundant genes, such as those involved in mitochondrial functions, may have limited influence on specific cellular capabilities. Consequently, researchers rely on comparative approaches, like differential gene expression (e.g., DESeq2 ^1^ and GSEA ^2^) and signature scoring (e.g., GSVA ^3^, Scanpy ^4^, Seurat ^5^), to quantify relative changes in expression, assuming this relative change is a proxy for functional importance. While insightful, these methods were not designed to generalize across experiments; each study’s unique design, cell type composition, and technical artifacts (like sequencing depth or batch effects) limit potential comparisons and complicate interpretation across datasets. Computational methods for normalization and dataset integration ^6–9^ have been developed to ameliorate such issues, but these approaches cannot be effectively applied to the thousands of publicly available single-cell transcriptomics experiments.

To enable scalable, robust, cross-dataset analyses and more objectively measure a gene’s functional importance, we developed the SIGnature (Scoring the Importance of Genes) framework, which adapts *attribution* methods from explainable artificial intelligence (XAI) and applies them to foundation models (FMs) trained on single-cell RNA-sequencing (scRNA-seq) data. Generally, attributions quantify the contribution of each input feature to a model’s prediction. For example, in image classification, the pixels with the highest attribution scores are most responsible for identifying an object, such as a dog in a backyard (**Figure 1a**). When applied to a scRNA-seq FM (**Figure 1b)**, attributions measure each gene’s influence on a cell’s position within the latent space of the model **(Figure 1c)**. If the FM encodes biological function, then genes with high attribution scores reflect their functional importance to a given cell. Indeed, we will demonstrate that attributions recover key markers of the cell’s identity or regulators of its specialized functions, and are more resistant to technical artifacts than normalized counts (**Figure 1d)**. Critically, attributions enable generalizable analyses across datasets because every cell is compared against the same standardized FM embedding.

**Figure 1:**
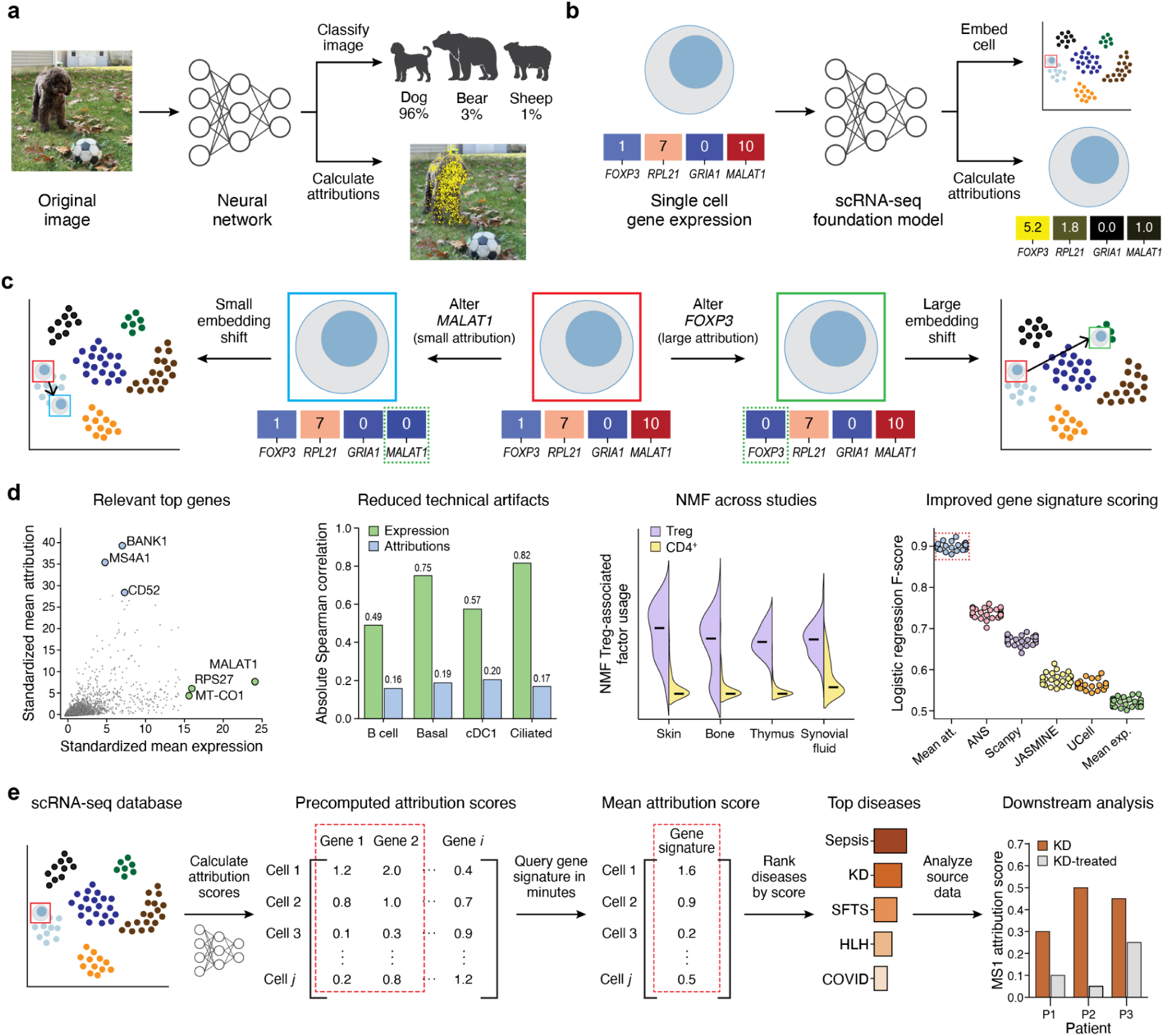
Gene importance is quantified through attributions. **a**. In image classification, attribution scores highlight key pixels (yellow) used by the model for the highest probability prediction (dog). **b**. In a scRNA-seq foundation model, attributions quantify each gene’s importance to the cell’s latent embedding position. Attribution values do not necessarily correlate with expression (*e.g*., *MALAT1* vs. *FOXP3*). **c**. Impact of gene expression changes on embedding: altering *MALAT1* (low attribution) has little effect; altering *FOXP3* (high attribution) causes major shifts. **d**. Key features of attributions in scRNA-seq analysis. Attributions enhance marker gene detection, reduce technical artifacts compared to log-normalized expression, and improve both cross-study NMF and gene signature scoring. **e**. SIGnature search workflow. Pre-computed attributions enable fast cell-level scoring for the genes in the signature (red box); atlas data reveal signature enrichment in cell types and diseases.

We demonstrate the utility of these attributions for biological discovery and drug development by rapidly querying established gene signatures across large scRNA-seq atlases and finding new associations between cell states, treatments, and diseases (**Figure 1e**). We focused on the MS1 gene program, a poorly understood myeloid phenotype linked to adverse outcomes in severe COVID-19 and sepsis ^10,11^. Using SIGnature to analyze over 400 diverse studies, we discovered activation of the MS1 signature in three previously unassociated inflammatory conditions: hemophagocytic lymphohistiocytosis (HLH), severe fever with thrombocytopenia syndrome (SFTS), and Kawasaki disease (KD). We experimentally validated the novel MS1 association with KD by showing that, similar to sepsis serum ^10^, KD patient serum can induce the MS1 phenotype *in vitro*. Further investigation into KD scRNA-seq data suggested MS1 cells were reduced after IVIG treatment, but *in vitro* testing could not confirm a direct association. These findings underscore SIGnature’s ability to reveal shared disease mechanisms and generate testable hypotheses through large-scale gene set scoring.

## Results

### SIGnature: Scoring the Importance of Genes using foundation model attributions

SIGnature is a framework for quantifying each gene’s importance to a single cell by calculating attributions against the embedding of a pre-trained FM. The first step is selecting a foundation model that meets two essential requirements: accepting a fixed set of genes as input and producing a cell-level embedding that forms a “biological universe” where distances are biologically meaningful. We consider a variety of pre-trained and fine-tuned FMs with diverse model architectures and loss functions: scFoundation ^12^, scGPT ^13^, SCimilarity ^14^, self-supervised models trained on scTab ^15,16^, and an scVI ^9^ model trained on the CellxGene Census data ^17^.

After selecting the pre-trained FM, we compute attributions using gradient-based explainability techniques – specifically integrated gradients (IG) ^18^, input x gradient (IxG) ^19^, and DeepLIFT (DL) ^20^ - because they scale well to the tens of thousands of features (i.e., genes) in scRNA-seq data (**Methods)**. To adapt these methods for multi-dimensional embeddings, a final summation layer is added to the pre-trained network ^21^ (**Methods**). This enables efficient transformation of a cell’s gene expression vector into an attribution vector of identical shape, where each gene’s score quantifies its contribution to the cell’s position within the model’s latent representation.

To better understand the utility of attributions, we used the SIGnature approach to calculate attributions on multiple datasets, using a variety of FMs and explainability methods (**Figure 2, Extended Data Fig. 2**). Benchmarking revealed that computation time is dependent on both the model and the method: calculating attributions with simpler MLP-based models is much faster than using transformer-based ones, and IG is the slowest of the three explainability methods (**Figure 2**). Furthermore, attributions from most models exhibit greater resistance to sequencing artifacts and successfully de-emphasize ribosomal genes relative to log-normalized expression. Additionally, attributions in all cases highlight mitotic genes in actively cycling cells, whereas attributions from models that incorporate cell type labels (i.e., SCimilarity and the fine-tuned SSL-scTab model) are most effective at increasing the relative importance of cell type marker genes (**Figure 2**). The boosting of biological signal is a key benefit of attributions; performing the same benchmarking with a shuffled gene order also reduces technical artifacts and de-emphasizes ribosomal genes, but it shows no enrichment of mitotic or cell type marker genes.

**Figure 2:**
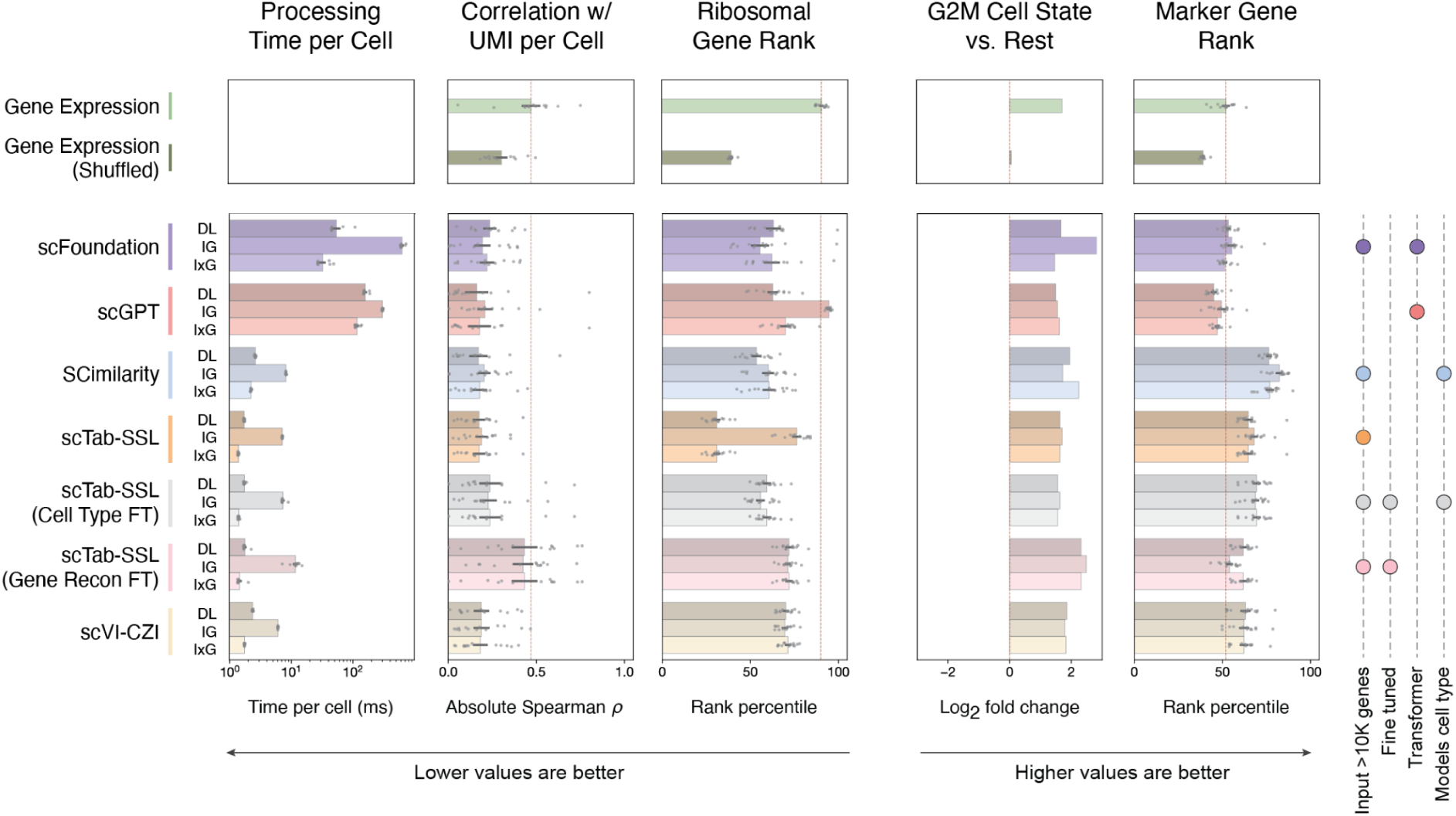
Benchmarking attributions across explainability methods and foundation models. The top section shows values for baselines: log-normalized gene expression using true and shuffled gene labels. Each row in the bottom section shows attribution values calculated using the indicated FM. This includes five pre-trained FMs (scFoundation ^12^, scGPT ^13^, SCimilarity ^14^, self-supervised model trained on scTab ^15,16^, and scVI ^9^ trained on CZI Census ^17^) and two fine-tuned versions of the scTab-SSL model (cell-typing and gene reconstruction ^15^). The right section details properties of each model. Center plots show the performance of three explainability methods: DeepLIFT (DL) ^20^, integrated gradients (IG) ^18^, and input x gradient (IxG) ^19^ across five evaluation criteria: 1) time to calculate attributions for a single cell (repeated 10 times), 2) correlation of marker gene attributions with cell UMI counts for each of thirteen blood cell types ^67^, 3) mean rank of ribosomal genes per blood cell type ^67^, 4) log_2_ fold change of G2M genes in mitotic fibroblasts vs. others ^69^, and 5) mean rank of marker genes per blood cell type ^67^. Lower values are better in the left three plots and higher values are better in the right two. Red dotted lines show the mean value for log-normalized expression, except in the G2M plot where the red line indicates the baseline value of 0 log2 fold change. Bar lengths display the mean value, error bars indicate the standard error, and grey dots display the individual values (each run for computation time and each cell type for blood dataset analysis).

These results establish attributions as a robust and valuable method for analyzing gene expression data. We therefore developed the SIGnature python package, enabling efficient attribution calculation using a suite of FMs and explainability methods. For our in-depth downstream analyses, we used the SIGnature package to investigate attributions calculated with the SCimilarity FM ^14^ and the integrated gradients method ^18^. This pair was selected because it leverages SCimilarity’s unique loss function, relatively small architecture, and large input space, alongside integrated gradients’ (**Figure 3a)** established balance of speed and performance. Henceforth, the term “attribution” in this manuscript refers specifically to scores calculated with this combination.

**Figure 3:**
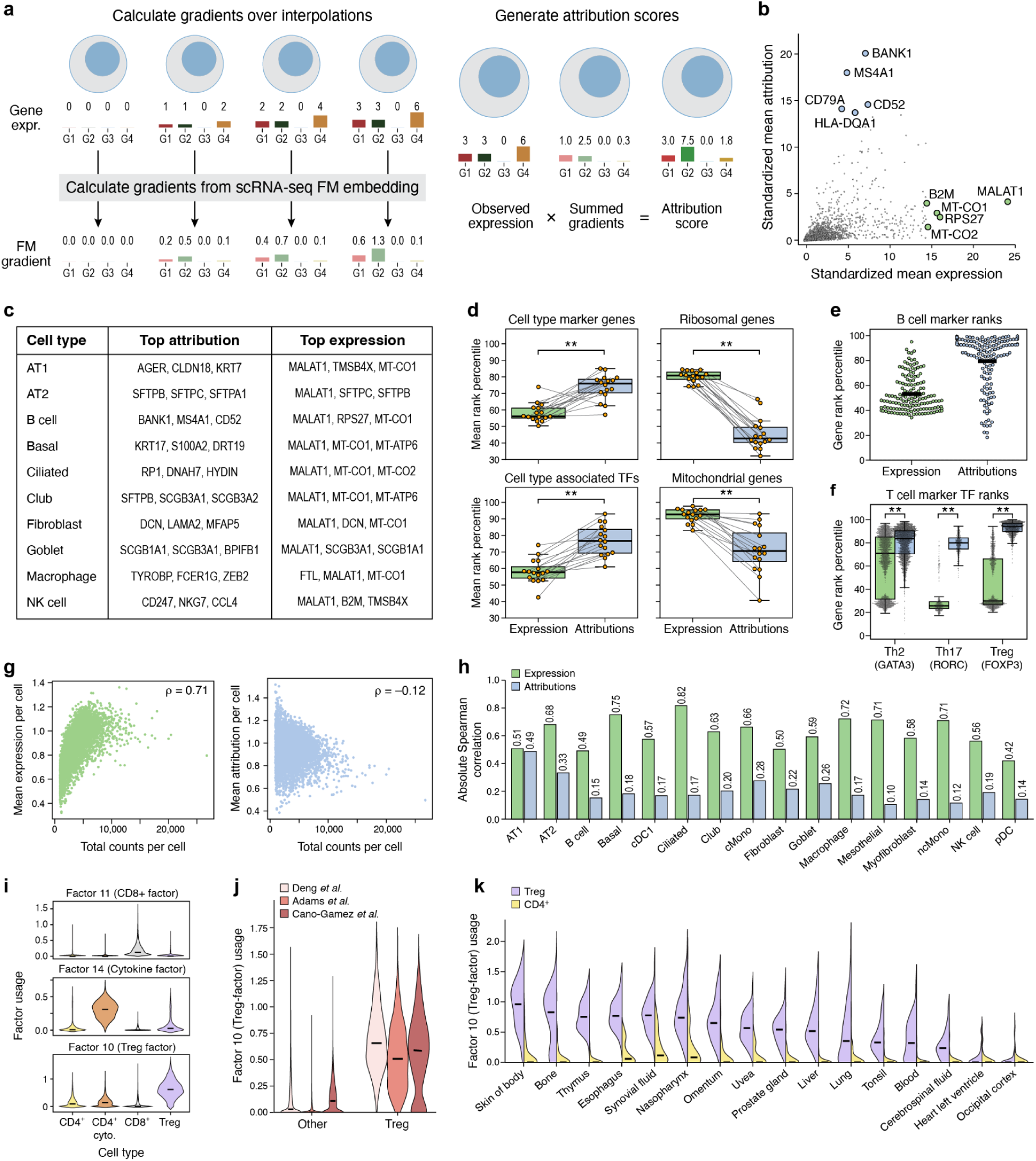
Attributions are biologically meaningful and useful for cross-study analyses. **a**. Attribution scores using a scRNA-seq FM are computed using gradients from interpolated expression profiles. **b**. Standardized mean expression (x-axis) and attribution (y-axis) for genes (dots) across B cells ^22^. Larger dots, labeled with corresponding genes, indicate top expression (green) and attribution (blue). **c**. Top genes by mean attribution (left) and expression (right) across lung cell types. **d**. Gene family rank comparison across lung cell types for four gene families (panel titles). Each dot shows a cell type’s average rank percentile for genes of that family; green and blue boxplots show expression and attribution quartiles, respectively. Same cell types are connected by thin black lines. (**p<0.01; Wilcoxon). **e**. Marker gene comparisons for B cells. Dots represent a marker gene’s average rank percentile across B cells. Thick black lines show mean values. **f**. In cytokine-treated T cells ^26^ (Th2, Th17, Treg), lineage TFs (*GATA3*, *RORC*, *FOXP3*) rank higher by attribution than expression (**p<0.01; Wilcoxon). **g**. Correlation of marker gene (dot) in non-classical monocytes (n=5,616): expression correlates strongly (ρ=0.71), attributions minimally (ρ=-0.12). **h**. Marker gene score correlations (y-axis) between expression (green) or attribution (blue) *vs.* sequencing depth across lung cell types (x-axis). Expression shows stronger correlation than attribution. **i**. NMF factor usage (y-axis) by cell type (x-axis). Factors shown correspond to CD8+ T cells (top), interferon response (middle), and Tregs (bottom). **j**. Treg factor usage (y-axis) across datasets used for NMF (legend). **k**. Treg NMF factor usage across tissues. Cross-study NMF factor applied to 100 predicted Tregs cells (purple) and 100 CD4+ T cells (yellow) not included in the original NMF training. Treg-specific factor scores are significantly elevated in predicted Tregs across all tissues (**p<0.01; paired t-test).

### Attribution scores capture biological importance and alleviate technical artifacts

Attribution scores reveal genes that are critical to a cell’s identity and functional role. We calculated attributions for B cells from a single lung scRNA-seq dataset ^22^ that was excluded from FM training, and found that the genes with the highest attributions, such as *BANK1* ^23^, *CD79A* ^24^ and *MS4A1* ^25^, have established associations with B cells **(Figure 3b)**. In contrast, genes with the highest expression counts (*e.g., MALAT1*, *RPS27*, and *MT-CO1*) are enriched for mitochondrial and ribosomal genes, as expected. This trend holds for many cell populations (**Figure 3c, Extended Data Fig. 3a**), indicating strong alignment between the genes the foundation model deems most important for a cell’s embedding and the known biological markers that define its identity and specialized functions. In fact, when ranking detected genes in each cell, cell type markers rank significantly higher in attribution scores than in expression levels (Wilcoxon test p < 0.01). Conversely, mitochondrial and ribosomal genes rank significantly higher in expression scores (Wilcoxon test p < 0.01) (**Figure 3d,e)**.

We then focused on transcription factors, which are crucial to cell functionality, but challenging to detect with scRNA-seq due to their low expression. Attributions enhance the ranks of TF markers across all tested lung cell types (**Figure 3d**). Additionally, when considering CD4+ T cells from another study ^26^, the lineage-defining TFs *GATA3*, *RORC*, and *FOXP3* show significantly higher within-cell attribution ranks than expression ranks in their corresponding T cell subsets (Th2, Th17, Tregs) (**Figure 3f**) (Wilcoxon test p < 0.01).

Finally, we show that attributions are robust to technical artifacts common in scRNA-seq data. Compared to expression, attribution scores of marker genes are less correlated with cDNA library complexity measures, such as total mRNA counts and the number of unique genes detected (**Figure 3g,h**, **Extended Data Fig. 3b**). Furthermore, attributions prove robust to random dropouts; when calculating attributions on B cells simulating dropout (50% of counts removed), the ranking of the top genes by attribution remains highly consistent (93% overlap) with the original, unmanipulated B cells (**Extended Data Fig. 3c**).

### Attributions facilitate cross-study gene program discovery

Given that attributions are biologically meaningful and robust within individual studies, we investigated whether they could enable cross-dataset analyses. Specifically, we sought to aggregate multiple datasets to enhance the identification of biologically relevant gene programs. For this task, we considered non-negative matrix factorization (NMF), which is commonly used on scRNA-seq data to identify gene programs ^27^, but is rarely applied to multi-experiment data due to the difficulty of separating biological signals from technical noise. We jointly analyzed T cells from three different experiments ^22,26,28^ by calculating attributions for every cell in each study and concatenating the three attribution matrices for NMF (**Methods**). Our analysis revealed interpretable gene programs, including factors specific to CD8+ T cells and cytokine-treated CD4+ cells (**Figure 3i, Extended Data Fig. 4a-d).** Notably, one factor highlights regulatory T cells (Tregs) across all three studies and has high loading scores for key markers *FOXP3* ^29^ and *IL2RA* ^30^ (**Figure 3j**). To assess the generalizability of this gene program, we used the learned weights to calculate usage scores for 3200 predicted T-cells from 16 tissues ^14^ (**Methods)** and observed significantly elevated scores in Tregs compared to other CD4+ T cells (**Figure 3k**, paired t-test, p < 0.01).

For this multi-study dataset, attribution-based NMF yielded more stable results across various random seeds, feature sets, and numbers of components than expression-based NMF (**Methods**). For instance, 90% of configurations using attributions produced a Treg-associated factor whose usage scores in Tregs were significantly elevated compared to the usage scores of other T cells across all three studies (t-test adj-p < 0.01) (**Methods)**. Additionally, 98% of those configurations consider key regulator, *FOXP3*, as a top 10 gene by loading weight. In contrast, performing the same analysis with normalized expression was less consistent: factors were often difficult to interpret (**Extended Data Fig. 4c,d)**, with only 58% of configurations yielding a Treg-associated factor, and only 4% of those cases identifying FOXP3 as a top 10 gene (**Extended Data Fig. 4e,f**).

We also observed that the quality of attribution-based NMF is influenced by the underlying FM’s representation. Here, we considered two multi-study datasets: the Human Lung Cell Atlas (HLCA) ^31^, composed of cell types that are heavily represented in SCimilarity’s training, and a single-cell atlas of the ocular surface ^32^, with corneal epithelial cells that are scarcely represented in SCimilarity. For the HLCA, attribution-derived factors are significantly more associated with cell type annotations than expression-derived factors and significantly less associated with the study of origin (Mann Whitney U adj-p < 0.01) (**Extended Data Fig. 5a-b)**. For example, one attribution-derived factor highlights ciliated cells and is consistently activated across multiple experiments and sequencing platforms (**Extended Data Fig. 5c,d)**. Analysis of the ocular data also showed that attribution-derived factors are less associated with study of origin than expression-based factors (Mann Whitney U adj-p < 0.01) (**Extended Data Fig. 5e)**. However, there is no significant increase in the fraction of cell-type associated factors for this dataset, suggesting the quality of attributions is limited by the foundation model’s representation of a given cell type (**Extended Data Fig. 5f)**.

These findings are also consistent across decomposition methods. Applying topic modeling to the expression data through Amortized LDA ^33,34^ or scETM ^35^ produces results similar to expression-based NMF (**Extended Data Fig. 5)**, with attributions generally boosting the cell type signal and reducing study-specific effects. Notably, attributions perform similarly to a supervised version of scETM topic modeling that incorporates batch labels. While supervised scETM performs slightly better on cell typing and equally well on reducing study-specific effects, this comparison highlights a key advantage of attributions as they achieve comparable results without requiring any new model training or parameter tuning.

In summary, performing NMF on attribution values is a powerful method for identifying robust, biologically meaningful gene programs that generalize across diverse datasets and technical platforms.

### Attributions enable signature scoring across scRNA-seq datasets

Having demonstrated the power of attributions in deriving meaningful, unsupervised gene programs across studies via NMF, we shifted to the challenge of signature scoring – quantifying the collective activity of known gene sets across cells from one or more studies ^4,5,36–39^. We found aggregating attribution scores across the genes within a gene set to be a highly effective approach for signature scoring. For example, in a single study of peripheral blood mononuclear cells (PBMCs) ^40^, the average attribution scores for a predefined list of each cell type’s marker genes were highest in the corresponding cell type **(Figure 4a, Extended Data Fig. 6a**). Furthermore, cell types with similar functional roles, such as the cytotoxically capable NK cells, CD8+ T cells, and δ-γ T cells, showed partially overlapping activation patterns. We also observed similar patterns in both single-cell and simulated spot-level spatial transcriptomics data ^41^: average attributions for cell type markers are elevated in appropriate cells and spots, and the magnitude of the attribution score often correlates with that cell type’s proportion in a given spot (**Methods, Extended Data Fig. 7)**.

**Figure 4:**
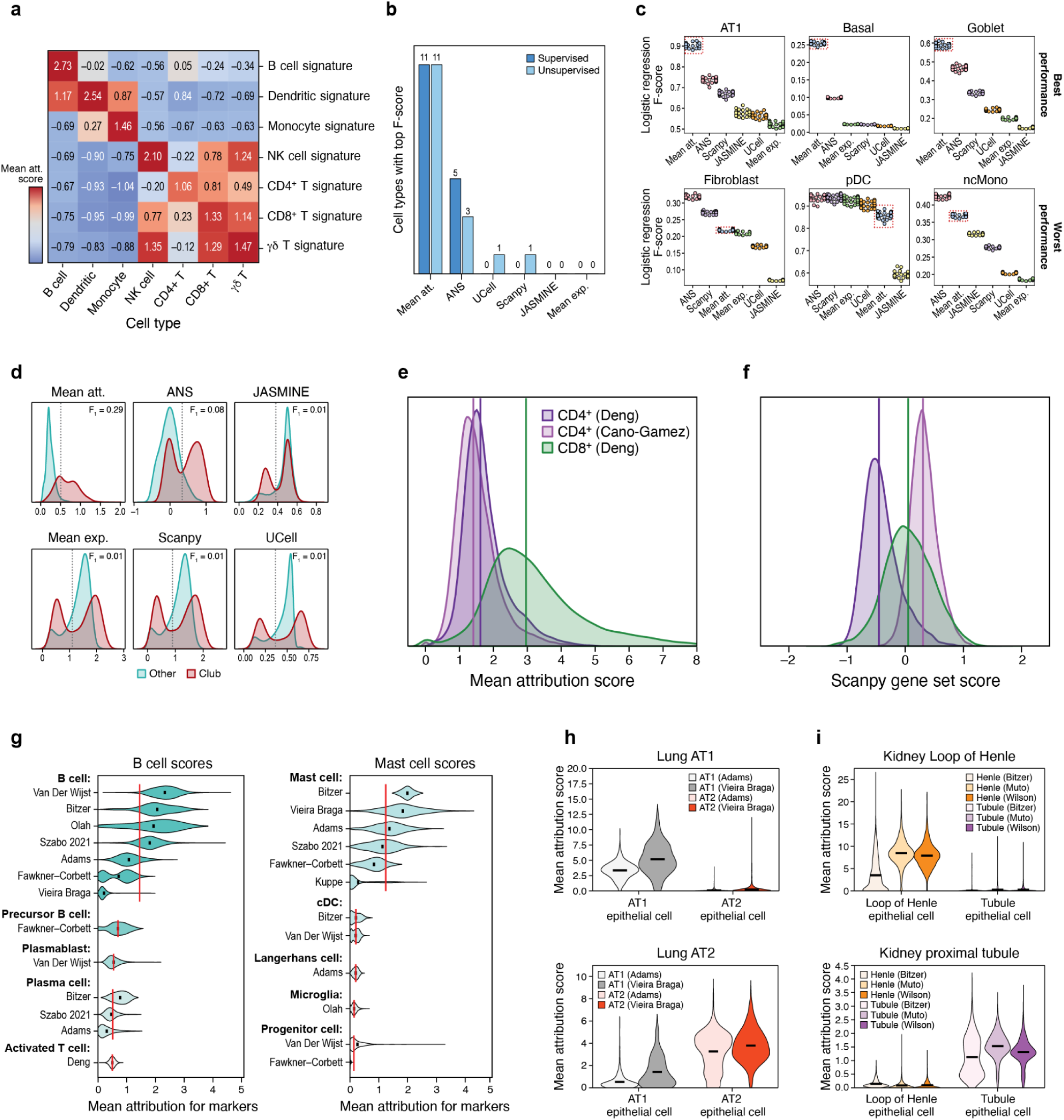
Average attributions measure gene set activation. **a.** Heatmap displaying standardized average attributions for marker genes (y-axis) in PBMC cell types (x-axis) (570,249 cells) ^40^. **b.** Classification performance for each scoring method (x-axis) across lung cell types (y-axis). **c.** Supervised classification F_1_-scores for top three best and worst performing cell types for attributions. Each dot represents the F_1_-score for one of 25 cross-validation runs. Red boxes highlight results for mean attributions. **d.** Distribution plots comparing club cell marker gene scores in club cells (maroon, n=2,492) *vs.* other lung cells (teal, n=279,302). Dashed lines indicate the Otsu threshold for determining F_1_-score. **e.** Density plots showing average attribution scores CD8+ T cell marker genes. CD8+ T cells from Deng *et al.* ^28^ (n=56,337, green) have higher scores compared to CD4+ T cells from Deng *et al.* ^28^ (n=16,794, purple) and Cano-Gomez *et al.* ^26^ (n=40,284, light purple). Medians indicated by vertical lines. **f.** Scanpy scoring of the same CD8+ gene signature assigns CD4+ T cells from Cano-Gomez *et al.* ^26^ higher scores than CD8+ T cells from Deng *et al.* ^28^ **g.** Marker gene attributions for B cells (left) and mast cells (right) across studies. Top five cell types by average study-level score are shown. Bolded text indicates cell type and y-axis label shows source study. X-axis indicates mean attribution score for marker genes. Black lines indicate mean values for each dataset, and red lines show the average of the study means for each cell type. **h.** Marker gene attribution scores for lung epithelial cell subtypes across multiple studies ^22,85^. **i.** Marker gene attribution scores for kidney epithelial cell types across multiple studies ^86–88^

We then explicitly compared mean attributions against established signature scoring methods in their ability to predict cell type. This comparison included background-based methods (ANS ^37^ and Scanpy ^4,39^) that score by comparing against dataset-specific background genes and rank-based methods (JASMINE ^38^ and UCell ^36^) that rely on within-cell ranking to quantify activation. Mean attributions achieved the highest F-score in 23/32 tests (**Figure 4b)** and was the best performing method across both supervised and unsupervised classification tasks (**Figure 4c,d, Extended Data Fig. 8**).

Average attribution scores are also robust across experiments. We first demonstrated this using two datasets that detected T cells ^26,28^. Considering a simple signature of CD8+ T cells (i.e., CD3 subunits [δ, ε, γ, ζ] and CD8 subunits [α, β]), the mean attribution scores clearly distinguish CD8+ T cells from CD4+ T cells across studies (**Figure 4e)**. In sharp contrast, applying Scanpy’s expression-based scoring to each study results in CD4+ T cells from Cano-Gamez *et al.* ^26^ scoring higher than the true CD8+ T cells from Deng *et al.* ^28^ (**Figure 4f**). This discrepancy highlights the inherent difficulty of applying dataset-specific scoring methods, such as Scanpy, across experiments and emphasizes the advantage of attribution-based scoring for cross-study comparisons. We extended this simple example by analyzing 1.2 million annotated cells from 15 experiments, consistently finding appropriate activation across studies in common immune cell types, like B cells and mast cells (**Figure 4g)**, and in diverse epithelial populations (**Figure 4h,i)**.

Furthermore, attributions can distinguish cell types not explicitly used during SCimilarity’s training, like the cDC1 and cDC2 subtypes of dendritic cells **Extended Data Fig. 6c,d)**, and can highlight cell states present across populations, such as cell cycle phase (**Extended Data Fig. 6e-g**). These results demonstrate that attribution scores are effective for detecting signature activation.

### Querying attributions across 22M cells reveals MS1-like state in multiple inflammatory conditions

Attributions from SIGnature also enable efficient atlas-level gene set querying, generating cell-level scores for 22 million cells in minutes, compared to hours or days with conventional methods ^42^. This is possible because attribution can be pre-computed for massive atlases and rapidly queried through simple mathematical operations. We leverage this capability in the following section to uncover a shared inflammatory cell state across diverse conditions, enabling the formulation of testable hypotheses.

To demonstrate the utility of gene set searching with SIGnature, we focus on the MS1 gene signature, which is activated in a poorly understood subpopulation of monocytes linked to adverse outcomes in sepsis and COVID-19 ^10,11^. We queried this signature of approximately 100 genes ^10,11^ across 412 human disease studies (22 million cells). Given that the signature is primarily enriched in myeloid populations (**Extended Data Fig. 9a**), all subsequent analyses focused exclusively on the 2.3 million predicted monocytes and macrophages. Cells with mean MS1 attribution scores in the top 10% were designated as MS1-like cells.

By calculating the prevalence of MS1-like monocytes/macrophages in each sample, we identified several diseases associated with MS1 activation (**Figure 5a, Extended Data Fig. 9b**). Our analysis recovered known biology, showing that the MS1 gene signature is most strongly associated with septic shock and more severe forms of COVID-19 ^43^ (**Extended Data Fig. 9c**). We also identified MS1-like cells in previously unassociated hyperinflammatory conditions: Kawasaki disease (KD) ^44^, severe fever with thrombocytopenia syndrome (SFTS) ^45^, and hemophagocytic lymphohistiocytosis (HLH) ^46^ (**Figure 5a**). These findings are robust against various hit thresholds (**Extended Data Fig. 9b**), suggesting that the MS1 signature is a disease biomarker for various inflammatory conditions.

**Figure 5:**
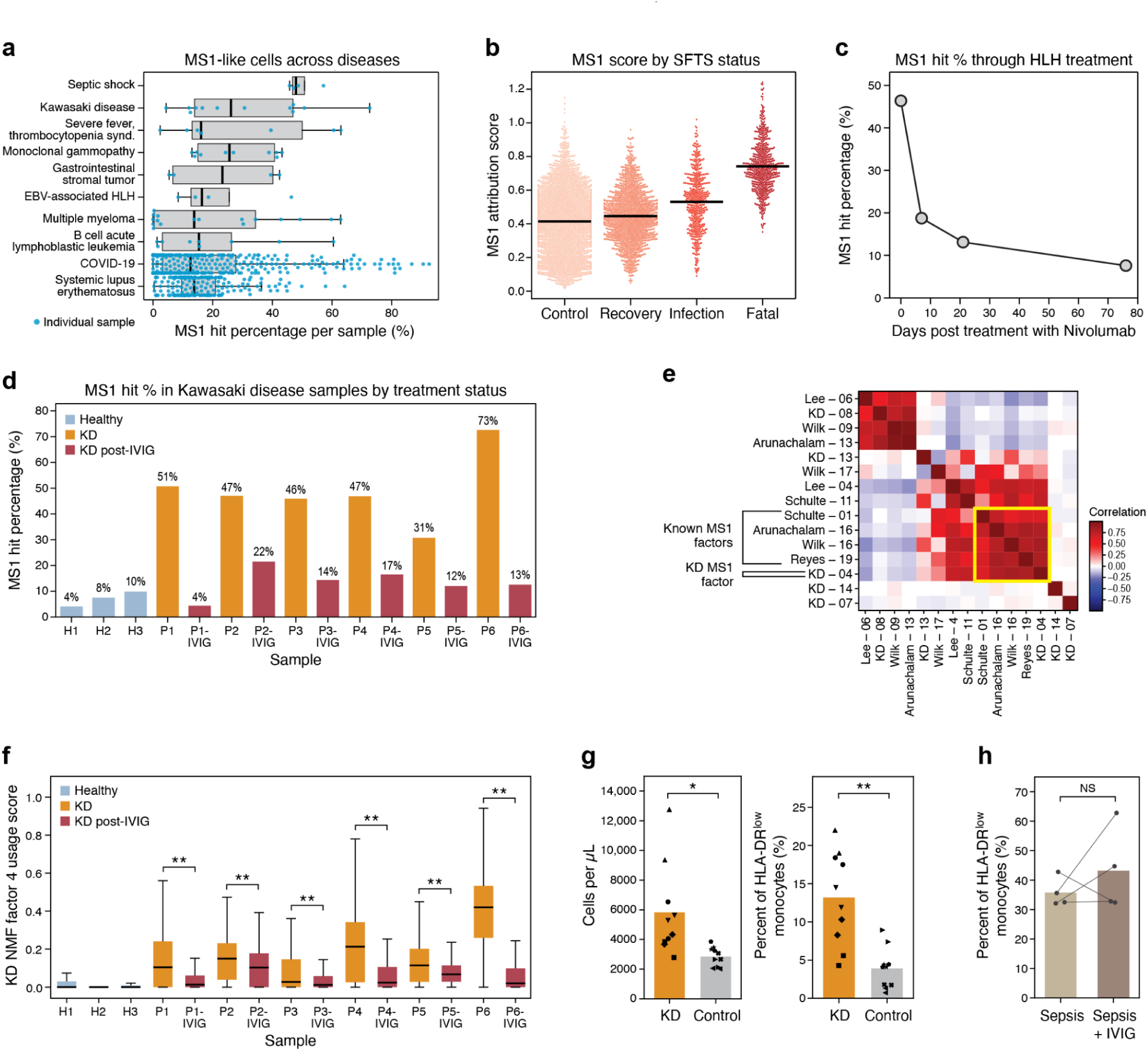
MS1 attribution scores are elevated in hyperinflammatory conditions. **a.** MS1 hit percentage by disease. The x-axis displays the percentage of monocytes/macrophages classified as MS1 hits per sample (dot), while the y-axis lists the top 10 diseases, ordered by mean sample hit percentage. Each dot represents a sample, and boxplots display quartiles. **b.** MS1 attribution scores across SFTS disease phases. Black lines indicate group averages. **c.** Decrease in MS1-like monocytes during Nivolumab treatment in an HLH patient. **d.** MS1 hit percentage in Kawasaki disease pre- and post-IVIG and in healthy controls (orange, red, and blue respectively). **e.** Clustering of NMF coefficients for monocyte gene programs. New KD programs cluster with established COVID-19 and sepsis programs ^10^; KD Factor 4 correlates with MS1 signatures (Arunachalam-16, Reyes-19, Schulte-01, Wilk-16) (**Extended Data** Fig. 10). **f.** KD Factor 4 scores by treatment status, with decreased usage post-IVIG (**p<0.01, Mann-Whitney U test). Boxplots show data quartiles, excluding outliers > 1.5 IQR; cell counts: H1 (n=442), H2 (n=200), H3 (n=355), P1 (n=418), P1-IVIG (n=1,767), P2 (n=576), P2-IVIG (n=144), P3 (n=586), P3-IVIG (n=181), P4 (n=640), P4-IVIG (n=121), P5 (n=1,222), P5-IVIG (n=390), P6 (n=833), P6-IVIG (n=407). **g.** KD serum induces emergency myelopoiesis and increases MS1-like cells. Left: Elevated monocyte concentration with KD serum compared to febrile controls (*p<0.05, mixed linear model). Right: Significant increase in HLA-DR^low^ monocytes (**p<0.01, mixed linear model). **h.** IgG has no effect on percentage of HLA-DR^low^ monocytes using sepsis serum (NS p=0.47, paired t-test). Thin lines connect the same patient between untreated (left) and treated (right) conditions.

To validate these novel findings, we re-analyzed the original datasets for these conditions, revealing new insights absent from the original publications. In the SFTS study ^47^, MS1 attributions correlate with disease severity: the scores are lowest in the control and recovery groups, elevated in infected samples, and highest in the most severe phenotype of fatal SFTS (**Figure 5b**, **Extended Data Fig. 9d**). Investigation of the HLH data ^48^ revealed the prevalence of MS1-like monocytes decreases throughout treatment with Nivolumab (**Figure 5c)**. Within the KD dataset ^49^ MS1-like cells were highly abundant in patients but decreased significantly (Wilcoxon p-value < 0.01) after administration of IVIG (**Figure 5d)**, the standard of care treatment in KD whose mechanism of action remains unclear ^44,49,50^. The association between MS1 attributions and KD was particularly striking because of its consistency across many patients and KD’s poorly understood disease etiology and treatment.

To confirm these KD findings were conserved in the expression data, we replicated the analysis used by Reyes *et al.* ^10^ to define the MS1 program and performed consensus NMF ^27^ on scRNA-seq data from KD monocytes (**Methods**). KD program 4 contains highly weighted loadings for core MS1 genes (e.g., *S100A9*, *RETN*) and clusters strongly with established MS1 factors from other diseases (**Figure 5f)**. Furthermore, Factor 4 is the only factor whose usage significantly decreases after IVIG treatment in all patients (Mann-Whitney U test, adj-p < 0.01). These results strongly support that KD MS1-like cells are conserved with those in sepsis/COVID-19, demonstrating that conventional expression-based analysis can reveal similar phenotypes when guided by attribution results.

### KD patient serum induces MS1 phenotype, validating a novel disease association

We then experimentally confirmed the MS1-like cells that SIGnature identified in KD were similar to those observed in sepsis ^10,11^. In their original studies ^10,11^, Reyes *et al.* showed that serum from patients with bacterial sepsis induces emergency myelopoiesis in hematopoietic stem and progenitor cells (HSPCs), resulting in high monocyte production and a significant increase in the proportion of MS1-like cells (CD45^+^, HLA-DRA^low^). Implementing a similar methodology, we found that KD patient serum also significantly increased monocyte production and the proportion of HLA-DRA^low^ monocytes compared to serum from febrile controls (**Figure 5g**). This suggests circulating factors in KD patient serum promote the expansion of this disease-associated monocyte population.

The successful *in vitro* induction of the MS1 phenotype led us to follow-up on another testable hypothesis generated from the SIGnature analysis. KD scRNA-seq data showed that MS1-like cells decrease after IVIG, which could either be a direct effect of IVIG on MS1 cells, or a secondary effect correlated with disease treatment. We tested whether IgG (the primary component of IVIG) affects MS1 induction rate, but observed no significant change in the proportion of MS1 monocytes (**Figure 5h**). These findings highlight the need for further investigation into the mechanisms by which IVIG modulates immune responses in KD.

## Discussion

The central contribution of our work is introducing foundation model attributions as a robust, standardized metric of gene importance. While explainable AI techniques have been applied in supervised transcriptomic classifiers ^51–53^, and methods like Cell2Sentence ^54^ have enhanced scRNA-seq interpretability, our approach uniquely leverages the latent knowledge encoded in FMs to derive a unified quantitative metric applicable to any cell. This framework boosts signal for crucial, low-expression genes, like TFs, and minimizes technical artifacts, enabling robust analyses like cross-study NMF. Critically, attributions can be precomputed, facilitating rapid gene set scoring across massive scRNA-seq atlases. These features enabled the development of the SIGnature package and directly facilitated the discovery of novel disease associations for the MS1 monocyte signature.

Despite its advantages, the SIGnature approach has several limitations. First, while the underlying FMs are explicitly trained to produce biologically meaningful cellular representations, attribution scores are primarily interpretative tools and may not be optimal for traditional scRNA-seq tasks, such as clustering. Second, attribution scores represent a complex transformation of gene expression values, making their units less interpretable compared to raw transcript abundances. Finally, the reliability of attribution outputs is inherently tied to the quality of the FM and the specific explainability method employed. For instance, our analysis showed that attribution-derived NMF provided significant benefit in cell types well-represented in the SCimilarity training data (lung tissue) but offered no meaningful advantage in poorly-represented data (ocular surface) (**Extended Data Fig. 5)**. Our benchmarking analysis further emphasizes that the original model’s complexity, training data, and loss function are crucial considerations, as they fundamentally influence both the attribution calculation and the resulting output values (**Figure 2)**.

Looking forward, we propose three key avenues for advancing this framework. First, methodological improvements in attribution calculation should be explored, including alternative approaches for computing attributions on multi-dimensional embeddings, gene masking strategies to enable perturbation-based explainability methods like SHAP ^55^, and aggregation methods to enhance robustness and consistency across multiple XAI approaches ^56^. Second, there is a need to develop specialized FMs tailored to specific biological applications. For example, FMs fine-tuned for targeted tasks could generate attributions optimized for deconvolution or the design of gene panels for spatial omics assays. Lastly, the SIGnature framework can be extended to additional modalities, including chromatin accessibility ^57^, spatial omics ^58^, and multi-modal data ^59^, enabling the calculation of feature importances across a diverse range of biological data types.

In summary, our manuscript demonstrates the significant potential of FM attributions as a standardized metric for gene importance, paving the way for further research to harness the knowledge embedded in genomic FMs for diverse and specialized biological applications.

## Methods

### Calculating attributions using explainable AI and scRNA-seq foundation models

Several methods exist for calculating feature attributions on deep learning models ^18,20,55,60,61^. These techniques, originally developed for supervised tasks like image classification, identify key input features that influence a model’s prediction. These methods can be broadly categorized into two groups: gradient-based and perturbation-based. Gradient-based methods like input x gradient (IxG) ^19^, integrated gradients (IG) ^18^, and DeepLIFT (DL) ^20^ quantify feature relevance by calculating gradients between the input features and the model output. In contrast, perturbation-based methods, like SHAP ^55^ and LIME ^62^, measure relevance by quantifying the effect on model outputs after altering sets of input features. This work focuses on gradient-based explainability methods due to their computational efficiency. Specifically, gradient-based approaches scale with the number of integration steps (m) rather than the number of input features (e.g., IG requires only O(m) backward passes per output). By contrast, perturbation-based approaches require repeated evaluations of the model on perturbed inputs; exact Shapley computation is exponential in the number of features (p), whereas KernelSHAP typically scales as O(p^2^) ^63,64^. Consequently, perturbation methods become impractical for high-dimensional settings such as single-cell transcriptomic data.

Single-cell RNA-seq foundation models are often trained in an unsupervised or semi-supervised manner to generate biologically meaningful high-dimensional embedding spaces. Even though the aforementioned explainability methods could, in principle, be used to interpret individual elements of these latent representations, this would require the embeddings to be *disentangled*—that is, for each latent dimension to correspond to a distinct and semantically meaningful biological factor. In practice, most existing models do not exhibit such disentanglement, and enforcing it would compromise model-agnostic applicability by imposing constraints on model design and training. Consequently, it remains challenging to determine how variations in gene expression influence the multiple latent variables within these representations. Furthermore, computing attribution scores for each latent dimension is computationally intensive, limiting the feasibility of these methods in such contexts. Here, building on our previous work ^21^, an efficient variation of IG for multi-dimensional embeddings was implemented using *Captum* ^65^. This approach adds a final layer to the original scRNA-seq FM, which calculates a weighted sum of each embedding dimension to yield a single output. This is more computationally efficient than calculating gradients against each latent variable independently and places greater emphasis on the dimensions that have large magnitudes in the cell’s embedding. Formally, let 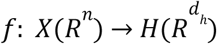 represent the scRNA-seq FM. The attribution score for the *i^th^* dimension of an input *X* is defined as follows:

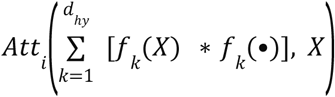

Building on this framework, three gradient-based attribution methods were applied to compute feature-level attributions against the final summation layer. “Input x Gradient”, an extension of the saliency method ^19^, calculates gradients for each feature against the model output and DeepLIFT ^20^ provides a more refined estimate by comparing each neuron’s activation to a reference activation and propagating contribution scores from the output back to the input. Finally, integrated gradients ^18^ computes attributions by interpolating between a baseline and the observed cell state (**Figure 2a**), calculating gradients for each interpolated input, and integrating these gradients to obtain the final attribution score. For both DeepLIFT and IG, a zero-vector was chosen as the baseline, following common practice in image-classification tasks, to ensure consistent comparisons across cells without dataset-specific tuning. For all three methods, the attribution values are then multiplied by the gene expression vector (preprocessed to be compatible with a given FM) and then re-normalized to a standard scale factor (i.e., 1000) to yield the final score for each gene, quantifying its influence on the cell’s representation in the embedding. Leveraging the SCimilarity model ^14^ and a batch size of 500 cells, this framework processes approximately 775 cells/s with IG and 3500 cells/s with DeepLIFT on a single L40S A100 GPU, illustrating the practical efficiency of these gradient-based approaches for large-scale single-cell data.

### Benchmarking attributions across scRNA-seq foundation models

Gradient-based measures of gene importance were implemented for five foundation models: SCimilarity ^14^ (v. 2023_01_rep0), scGPT ^13^ (whole-human model), scFoundation ^12^, an SCVI-based ^9^ model trained on CELLxGENE data ^17^ (2023-12-15-scvi-homo-sapiens), and a self-supervised model implemented with random masking on the scTab dataset ^15,16^. Additionally, these approaches were applied to two fine-tuned models ^15^ that were originally trained on the scTab dataset and then re-trained using self-supervised learning on Tabula Sapiens dataset ^66^ with the goals of cell type classification (CT) or gene reconstruction (GR) respectively. Attribution scores were calculated using the *Captum* ^65^ library to implement three aforementioned gradient-based methods (DL, IG, and IxG). For each FM, attributions were calculated against the primary embedding layer. For the fine-tuned classification model, attributions were calculated against the second to last network layer.

The attributions collected from these models were compared against each other on multiple tasks assessing their computational performance, biological relevance, and resistance to technical artifacts. Two additional baselines were included: log-normalized expression and log-normalized expression using shuffled gene labels, where **Figure 2** shows the average result from 25 shufflings. Most tasks used a downsampled version of the Wells *et al*. ^67^ dataset (n=2,943 cells), sampling at most 15 cells from each donor for each of the 14 cell types that have matching gene sets from CIBERSORT ^68^ (**Supplementary Table 2)**. Additionally, one task considered fibroblasts annotated with cell cycle state from Riba *et al.* ^69^.

Speed was assessed by measuring the time required to generate gene attributions for a single, pre-processed cell on an L40S GPU. This procedure was repeated for 10 unique cells and summarized in **Figure 2**. It should be noted that model performance can be further optimized by processing multiple cells concurrently on the GPU, a strategy that particularly benefits smaller models as more cells can be loaded onto the GPU at once.

To evaluate robustness, the mean value (either attribution or log-normalized expression) of established marker genes from CIBERSORT ^70^ was calculated for each cell type (**Supplementary Table 2**), and its absolute Spearman correlation with the cell’s total counts was determined. Two marker gene analyses were also performed using within-cell ranking. Non-zero gene values rank-normalized from 0 to 100 in ascending order using the scipy “rankdata” method ^71^ with default parameters. For each cell type, the average rank percentile was calculated for each gene in every cell in which it was detected. The mean value for all genes in a given gene set (e.g., ribosomal genes or cell type markers) is reported in **Figure 2**. Lastly, cell cycle analysis was performed using the G2/M phase genes from Tirosh *et al.* ^48^. A G2/M score was calculated for each cell as the mean value of these genes, and **Figure 2** reports the log_2_-fold change of this score between G2M-annotated cells relative to all other cells.

Each model uses a unique fixed set of genes, and attributions can only be calculated for those genes included in that specific model’s input space and the corresponding dataset of interest. **Figure 2** shows results including all genes that each model is capable of utilizing, besides scGPT which used 3000 highly variable genes because of memory limitations on the GPU. **Extended Data Fig. 2** shows the same analysis, but only using the included in the input space for all five foundation models (3497 and 1395 for the Wells *et al.* ^67^ and Riba *et al.* ^69^ studies respectively).

### Datasets and gene sets

24 publically available datasets were used for focused analyses in this manuscript. These studies are described in **Supplementary Table 1** and concisely summarized in **Extended Data Fig. 1**. Additionally, many tests in this manuscript used established gene sets associated with particular cell types or molecular phenotypes. **Supplementary Table 2** lists the genes included in each gene set and the cell type annotations with which they were associated.

### Plotting and statistics

Plotting was performed in python using *seaborn* (v. 0.13.2) ^72^ and *matplotlib* (v. 3.9.0) ^73^. Statistical tests were implemented using *scipy* (v. 1.14.0) ^71^ and *statsmodels* (v. 0.14.4) ^74^ .

### Top attributions per cell type

The Adams *et al.* lung dataset ^22^ includes the 16 cell types that have a corresponding lung marker gene set from MSigDB ^2,75^. For each of these 16 cell types, the average log-normalized gene expression and the average attribution values were calculated for every gene. **Figure 3b** shows a scatterplot of gene values in B cells, where the x-axis indicates the standardized average expression values and the y-axis shows the standardized average attribution values. **Figure 3c** shows the top 5 ranking genes by average attribution and expression for each cell type ^76^.

The average rank analysis (**Figure 3d-e**) was the same as benchmarking, with non-zero gene values rank-normalized from 0 to 100 in ascending order using the “rankdata” method from *scipy* ^71^ and taking the average rank for tied values. For each cell type, the average rank percentile was calculated for each gene in every cell it was detected. **Figure 3d** shows the summary for four gene families: lung cell type marker genes from MSigDB ^2,75^, ribosomal genes annotated from KEGG ^77,78^, mitochondrial genes (starting with MT-), and marker genes from MSigDB that are also transcription factors ^79^. The values for each individual cell type and gene family are shown in **Extended Data Fig. 3a.**

### Correlation analysis

For each of the 16 cell types analyzed from Adams et al. ^22^, the average log-normalized expression and attribution scores were collected for marker genes in each cell. The absolute Spearman correlation was then calculated between these values and three scRNA-seq QC metrics: number of UMIs, number of unique genes, and percent of UMIs from mitochondrial genes. **Figure 3g** shows an example comparing the total UMI in each cell with the average counts and attributions for non-classical monocytes. The results across all cell types are summarized in **Figure 3h** and **Extended Data Fig. 3b**.

### Robustness analysis

Each B cell from Adams *et al.* ^22^ was downsampled by randomly removing X% of the counts in each cell for the percentages: 0%, 10%, 20%, 30%, 40%, 50%, 60%, 70%, 80%, 90%. These downsampled datasets were then log-normalized and used to calculate attribution scores. For each downsampling percentage, the average attribution score for each gene was calculated across all B cells and ranked in descending order. The top scoring genes were robust to downsampling, with 93 of the top 100 genes remaining in the top 100, even after 50% of the counts had been removed (**Extended Data Fig. 3c**).

### Cross-study non-negative matrix factorization and topic modeling

Non-negative matrix factorization (NMF) was implemented using *scikit-learn* (v. 1.5.0) ^80^ on author-annotated T cells from three studies: Adams *et al.* ^22^, Cano-Gamez *et al*. ^26^, and Deng *et al.* ^28^. Analysis included 14,056 genes detected across all three studies. NMF was performed 200 times for both expression and attributions, varying three parameters: number of factors (5, 10, 15, 20, 25), random state (114-133), and feature set (all 14,056 genes or 2,591 highly variable genes).

For each run, a "Treg-associated factor" was identified by calculating t-statistics between Tregs and non-Tregs across all nine possible within-study and cross-study comparisons (*scipy* “ttest_ind”, alternative = “greater”). The factor with the highest minimum t-statistic across these nine comparisons was selected as the potential Treg-associated factor. The Treg-associated factor was considered to robustly highlight Tregs if that the worst performing Treg vs. non-Treg comparison was significant, using a Bonferroni-adjusted p-value of 2.7E-6 (alpha=0.01 divided by 3600 tests: 400 trials, each with 9 comparisons). **Figures 2i-k** show results from a specific parameter set (10 factors, all genes, random_state=114), though similar trends were observed across most parameter combinations.

Generalization analysis was performed using 3,200 cells from the SCimilarity database ^14^ that were not used for model training or validation. Sixteen tissues were selected because they contain at least 100 Tregs and 100 CD4+ T cells, as predicted by SCimilarity (nn_prediction_dist < 0.02). The NMF models trained on the original T cell cohort were applied to this independent dataset to assess generalization (**Extended Data Fig. 4e-f**).

Additional validation was performed on two independent studies downloaded from CZI: the Human Lung Cell Atlas (HLCA) ^31^ and a single-cell atlas of the ocular surface ^32^. For the HLCA, the dataset was downsampled to include at most five cells of each cell type (annotation level 5) from each sample not included in the SCimilarity training set (n=34,439). Genes were filtered to those present in at least 0.5% of cells (n=15,133). For the ocular surface atlas, all epithelial cells (n=12,990) were included, and genes were similarly filtered to those detected in at least 0.5% of cells (n=14,816). Attributions for both datasets were then calculated using the SCimilarity and IG. NMF was performed using the same 200 parameter settings as the T cell analysis. Additionally, topic modeling was performed using the scETM ^35^ and the “AmortizedLDA” method *scvi-tools* ^33^. scETM was run for 500 epochs with and without batch supervision using the “enable_batch_bias” parameter. Otherwise, both scETM and AmortizedLDA were implemented for each of the same k values (5, 10, 15, 20, 25) on both the full dataset and highly variable genes using the default parameters and a seed of 114.

The *statsmodels* ^74^ package was used to fit a linear model (FACTOR_SCORE ∼ C(cell_type)) to test if the transformed NMF or LDA values could be explained by cell type. The “ann_level_3” and “celltype_legency” columns were used for cell type annotations in the HLCA and ocular datasets respectively. For each parameter set, factors or topics with an adjusted R-squared value greater than 0.25 were considered as cell-type associated. This same analysis was then repeated to determine if factor usage was predicted by the cell’s study of origin. The results are presented in **Extended Data Fig. 5**.

### Spatial Transcriptomics Analysis

Slide-TAG spatial transcriptomics analysis used tonsil data from Russell *et al.* ^41^. Cell type labels were combined into 5 large immune cell categories: Bcell (B_germinal_center, B_naive, and B_memory), Tcell (T_CD4, T_follicular_helper, T_CD8, T_double_neg), Plasma (plasma), NK (NK), and Myeloid (mDC, myeloid, pDC). Simulated Visium spots were created by generating a grid of 25 micrometer spots and aggregating counts from cells within the border of each spot, converting 3199 cells into 625 spots. Attributions were then calculated on log-normalized counts from the single cells and the simulated spots using SCimilarity and integrated gradients. Downstream analysis is shown in **Extended Data Fig. 7**. Regression plots in **Extended Data Fig. 7d** were created with the “regplot” function from *seaborn* ^72^ using the default parameters.

### Dataset specificity and scanpy scoring

The “score_genes” function from *Scanpy* was implemented on two datasets: Deng *et al.* ^28^ and Cano-Gomez *et al.* ^26^. The gene set queried included all isoforms of CD3 (*CD3D*, *CD3E*, *CD3G*, and *CD247*) and of CD8 (*CD8A* and *CD8B*). Density plots were created using the “kde_plot” function in *seaborn* ^72^ with standard parameters.

Additionally, **Extended Data Fig. 6a** demonstrates that *Scanpy*’s “score_genes” function produces dataset-dependent results. When applied to cDC2 marker genes ^81^ across the same cells in different contexts (full Adams *et al.* ^22^ dataset versus a subset containing only dendritic cells and mast cells), the scores shift systematically, highlighting the method’s sensitivity to background cell composition.

### Implementing established gene set scoring methods

Gene set activation was evaluated by comparing mean attributions against five existing methods: Adjusted Neighborhood Scoring (ANS) ^37^, JASMINE ^38^, Mean Expression, Scanpy ^4,39^, and UCell ^36^. For all methods, data were log-normalized in Scanpy (v. 1.9.2) with a target sum of 10,000. Scores were calculated separately for each dataset to accommodate methods using background distributions.

Mean Expression averages log-normalized expression for genes of interest. Scanpy scoring uses the "score_genes" function with default parameters, which is based on the Tirosh *et al.* ^5,39^ approach, comparing query gene expression against background genes with similar expression levels. ANS ^37^ also uses control genes but identifies backgrounds through gene expression neighbor graphs rather than expression buckets. UCell ^36^ and JASMINE^38^ are both rank-based gene set scoring approaches; UCell calculates Mann-Whitney U statistics from expression ranks ^36^, while JASMINE computes likelihood or odds ratios from within-cell gene ranks ^38^. ANS, UCell, and JASMINE were implemented using the ANS_signature_scoring package ^37^, with ANS using a control_size of 50 and JASMINE using the “likelihood” model.

### Gene set scoring across scimilarity validation dataset

The gene sets used in this study are summarized in **Supplementary Table 2.** Immune gene sets from CellMarker 2.0 ^82^ and CIBERSORT ^70^ were applied to PBMCs from Van Der Wijst et al.^40^. Seven cell types with matching gene sets from both sources were analyzed. (**Extended Data Fig. 6b)**. For each cell, consensus scores were created by averaging marker gene scores from both sources for the same cell type. These consensus scores were then averaged by annotated cell type identity and displayed in **Figure 4a**.

For analysis across all 15 SCimilarity datasets, B cells and mast cells were considered as these populations appeared in the most unique studies (7 and 6 respectively). CIBERSORT ^70^ signatures were used to query these immune cell populations across datasets (**Figure 4d)**. Lung and kidney epithelial cell signatures were collected from CellMarker 2.0 ^82^ (**Figure 4e,f)** and dendritic cells were assessed using both CellMarker 2.0 and from Table 1 of Collin *et al.* ^81^ (**Extended Data Fig. 6c,d)**. **Extended Data Fig. 7e-g** was generated using cell cycle gene sets for S and G2/M phases from Tirosh *et al.* ^48^ to show that attributions can identify these cell states across cell types, including annotated fibroblasts ^69^.

### Supervised and unsupervised cell type prediction

Cell type classification performance was evaluated using six gene set scoring methods: ANS ^37^, JASMINE ^38^, Mean Attribution, Mean Expression, Scanpy ^4,39^, and UCell ^36^. For each cell type from Adams *et al.* dataset ^22^ and each method, marker gene set scores (**Supplementary Table 2**) were used to train univariate logistic regression classifiers using LogisticRegressionCV from scikit-learn (v. 1.2.0) ^80^ with balanced class weights and a random state of 114. Performance was assessed across 25 independent train/test splits (80%/20%) using F-score to account for class imbalance

Unsupervised classification employed Otsu’s multi-thresholding method ^83^, using the “threshold_multiotsu” function from scikit-image (v. 0.19.3)^84^. For each cell type, marker gene set scores were calculated for each cell and optimal thresholds determined. Multiple threshold numbers (n=1,2,3) were evaluated, with optimal N selected by maximizing Cohen’s d effect size between the top group and second-highest group, ensuring robust separation between populations. This approach provides an objective threshold selection method when true labels are unavailable. The cells above the chosen threshold were considered “hits” and used to calculate an F_1_-score against the true labels. Classification results are shown in **Extended Data Fig. 8**.

### MS1 full dataset queries

The MS1 phenotype in monocytes was established through NMF analysis by Reyes *et al.* (2021) ^10^. Attributions were queried across the SCimilarity cell database ^14^ using the top 99 genes associated with this factor to identify other cells activated for this phenotype. This search queries 22M cells across 412 datasets, which was then reduced to 2.3 million cells from 244 studies that were confidently predicted as monocytes or macrophages (SCimilarity prediction_nn_dist < 0.02). For sample-level hit analysis (**Figure 4a**), samples were considered if they contained at least 25 monocytes/macrophages and were from a disease with at least 3 unique samples. Hits were called as the top 10% of monocytes or macrophages by MS1 score and the percentage of hits was calculated for each sample (**Figure 4a**). Other thresholds (5%, 15%, and 20%) were also tested and showed similar top diseases (**Extended Data. Fig. 9b**).

### MS1 Cells in Individual Datasets

The MS1+ hits from the SCimilarity database were analyzed in the context of their respective datasets. Individual analyses were performed for multiple diseases using metadata from the original study: COVID-19 from Stephenson *et al.* ^43^, HLH from Liu *et al.* ^48^, SFTS from Park *et al.* ^47^, and KD from Wang *et al.* ^49^.

Consensus NMF was run on scRNA-seq data from the 8282 monocytes/macrophages from Wang *et al.* ^49^ using the *cNMF* python package ^27^ in order to compare results against COVID-19 and sepsis samples processed in Reyes *et al.* ^10^. The expression data from these cells were run through the *cNMF* package using the top 3000 variables genes, considering k values from 5 to 20, and repeating each trial 25 times. The optimal k was chosen as 14 based on the balance between high stability and relatively low error (**Extended Data Fig. 10a**).

The standardized gene loadings from the KD monocytes were compared against the standardized gene loadings from Reyes *et al.* ^10^ by computing pairwise Pearson correlation coefficients. These values were clustered using the *seaborn* clustermap function with the “correlation” metric ^72^. **Figure 5e** shows a subset of this clustered matrix, with the entire clustermap displayed in **Extended Data Fig. 10b**. For each patient, a Mann-Whitney U test was performed comparing the usage values for cells collected before IVIG against those collected after IVIG.

### MS1 Induction Experiments

Differentiation of myeloid cells from CD34+ bone marrow hematopoietic stem and progenitor cells (HSPCs) was performed as previously described ^10^. HSPCs cells were cultured in SFEM II with 75 nM StemRegenin 1, 3.5 nM UM729, 1X CC110 (StemCell Technologies), and 1X penicillin-streptomycin (Gibco). To initiate differentiation, cells were cultured in the same medium supplemented with 20% pooled healthy human serum (SeraCare) or serum from diseased patients. Serum samples from patients with multiple myeloma were obtained from BioIVT.

To assess MS1 induction, cells were stained with the following panel: CD14-FITC (clone M5E2), CD15-APC (clone W6D3), CD11b-AF700 (clone ICRF44), CD34-BV650 (clone 561), HLA-DR-PE/Cy7 (clone L243) (BioLegend) and IL1R2-PE (clone 34141; Thermo). Cells were resuspended in FACS buffer with 5% CountBright beads (Invitrogen) to allow determination of absolute counts during analysis. Flow cytometry data were acquired on an LSR Fortessa (BD Biosciences) and analyzed using FlowJo v10.10. For analysis of HLA^low^ monocytes, the average HLA intensity value was calculated for internal controls and then for each sample, the percentage of monocytes with HLA intensity below that value was calculated. Mixed linear analysis was performed using the *statsmodels* ^74^ package in python.

## Data Availability

The datasets used in this manuscript are summarized in **Supplementary Table 1** and **Extended Data Fig. 1** Precomputed attribution scores for the SIGnature package can be found on Zenodo (https://zenodo.org/communities/signature).

## Code Availability

The SIGnature package is available on Github (https://github.com/Genentech/SIGnature). Model files necessary for generating new attributions can be found on Zenodo (https://zenodo.org/communities/signature).

## Acknowledgements & Funding

We thank J. Collier, Y. Lee, A. Andersson, B. Chen, S. Malek, and M. Crow for their suggestions on the manuscript and L. Gaffney for her assistance in figure generation. We also thank S. de Ferranti and K. Bonello for their help in biological interpretation and sample procurement. MBS is supported by the Samara Jan Turkel Clinical Center for Pediatric Autoimmune Disease.

## Author Contributions

MPG, GH, and TB designed the study; MPG developed the method with ND, GH, AT, HCB, TK, TB, and EH; Model implementation was performed by MPG, ND, TK, and EH; Python API was developed by TK, with help from MPG; MPG performed the data analysis with guidance from GH, TB, AB, AT, HCB, MR, and SBK; MPG and MR conceived of the biological validation, with input from SBK, PYL, MBFS, and JWN; PYL, MBFS, and JWN provided advice on biological interpretation and coordinated sample procurement; MR performed the experimental validation. MPG, GH, and TB wrote the manuscript with input from EH and HCB; All authors reviewed the manuscript.

