## Extended Data Figures for "Scoring gene importance by interpreting single-cell foundation models"

| Study | FM Benchmark | Cell Type Scoring | Scanpy Gene Set Scoring | TCell NMF | Lung NMF | Eye NMF | Spatial Analysis | Cell Type Classifiers | Marker Gene Correlation | Cell Cycle Analysis | MS1 analysis | Scimilarity Training |
|---|---|---|---|---|---|---|---|---|---|---|---|---|
| Riba *et al.* (2022) | Y | | | | | | | | | Y | | |
| Wells *et al.* (2025) | Y | | | | | | | | | | | |
| Szabo et al. (2019) | | Y | | | | | | | | | | |
| Szabo et al. (2021) | | Y | | | | | | | | | | |
| Ravindra et al. (2021) | | Y | | | | | | | | | | |
| Fawkner-Corbett et al. (2021) | | Y | | | | | | | | | | |
| Van Der Wijst et al. (2021) | | Y | | | | | | | | | | |
| Kuppe et al. (2022) | | Y | | | | | | | | | | |
| Muto et al. (2021) | | Y | | | | | | | | | | |
| Deng et al. (2020) | | Y | Y | Y | | | | | | | | |
| Vieira Braga et al. (2019) | | Y | | | | | | | | | | |
| Cano-Gamez et al. (2020) | | Y | Y | Y | | | | | | | | |
| Adams et al. (2020) | | Y | Y | Y | | | | Y | Y | | | |
| Henry et al. (2018) | | Y | | | | | | | | | | |
| Olah et al. (2020) | | Y | | | | | | | | | | |
| Bitzer (2022) | | Y | | | | | | | | | | |
| Wilson et al. (2022) | | Y | | | | | | | | | | |
| Sikkema et al (2023) | | | | | Y | | | | | | | |
| Chen (2025) | | | | | | Y | | | | | | |
| Russell *et al.* (2023) | | | | | | | Y | | | | | |
| Stephenson et al (2021) | | | | | | | | | | | Y | Y |
| Park et al (2021) | | | | | | | | | | | Y | |
| Liu et al (2020) | | | | | | | | | | | Y | |
| Wang et al (2021) | | | | | | | | | | | Y | |

**Extended Data Fig. 1: Dataset Summary**
Every column indicates an analysis performed in the manuscript. Each row represents a study and shows the specific analyses in which it was used (marked with a "Y").

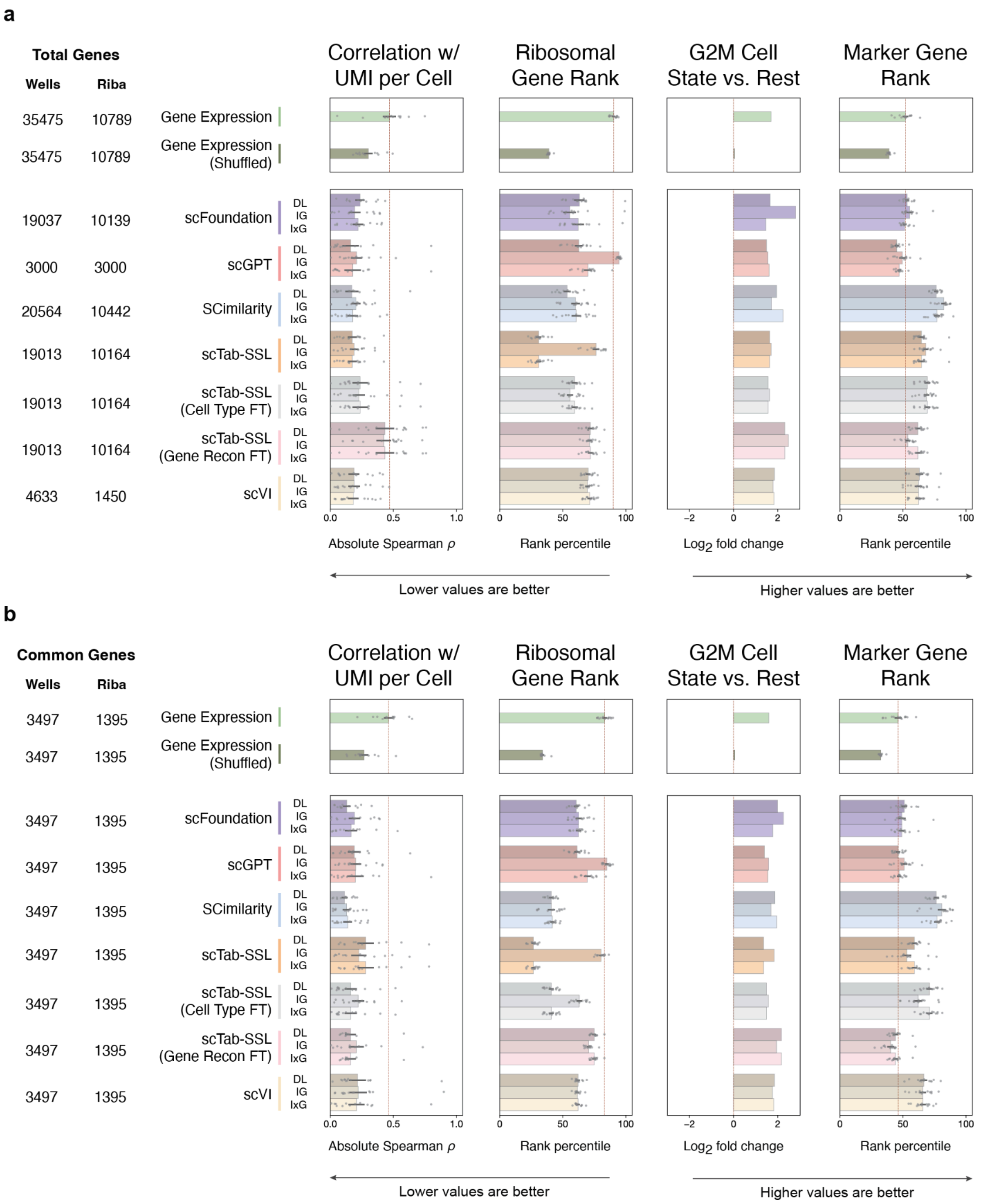


**<u>Extended Data Fig. 2</u>: Benchmarking models and explainability methods across feature sets**
**a.** benchmarking using all genes. **b.** benchmarking using common genes. Each section a shows the same data as **Figure 2,** except the legend on the left side now indicates the number of genes common between each model's input space and the dataset of interest: Riba *et al.* [1] fibroblast study used for column 3 and the Wells *et al.* [2] blood cell dataset used for the other analyses. The results are similar whether considering all genes (a) or common genes (b).

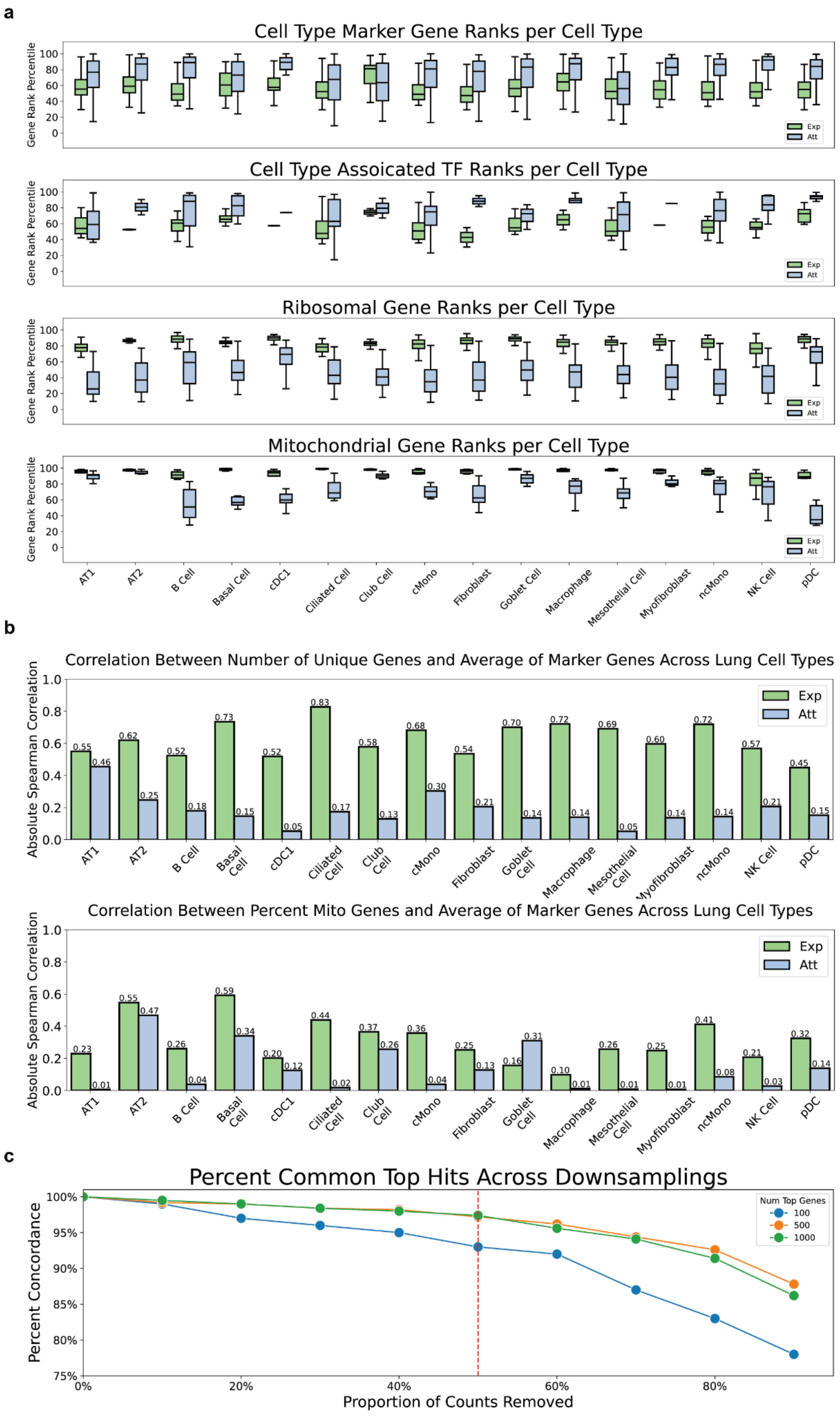
a
Cell Type Marker Gene Ranks per Cell Type
Cell Type Assoicated TF Ranks per Cell Type
Ribosomal Gene Ranks per Cell Type
Mitochondrial Gene Ranks per Cell Type
Gene Rank Percentile
Exp
Att
AT1
AT2
B Cell
Basal Cell
cDC1
Ciliated Cell
Club Cell
cMono
Fibroblast
Goblet Cell
Macrophage
Mesothelial Cell
Myofibroblast
ncMono
NK Cell
pDC
b
Correlation Between Number of Unique Genes and Average of Marker Genes Across Lung Cell Types
Correlation Between Percent Mito Genes and Average of Marker Genes Across Lung Cell Types
Absolute Spearman Correlation
c
Percent Common Top Hits Across Downsamplings
Percent Concordance
Proportion of Counts Removed
Num Top Genes
100
500
1000

**Extended Data Fig. 3: Attributions in lung cell types**

**a.** Expression and attribution ranks for a given gene family in a cell type of interest from Adams *et al* [3]. The x-axis displays the cell type and the y-axis shows the percentile. Each box indicates the data quartiles with outliers beyond 1.5X the IQR not shown. The genes of interest are indicated in the plot title. **b.** Correlation between values of interest and number of genes detected per cell. The x-axis is the cell type of interest and is split between the mean expression (green) and mean attribution (blue). The y-axis indicates the magnitude of the spearman correlation between the given scoring value and the sequencing metric of interest per cell for all cells in the cell type indicated by the x-axis label. **c.** plot showing the same values for the percentage of counts from mitochondrial genes.

**a**

Attribution-based NMF Top Genes per Factor (All Genes)

| | NMF-1 | NMF-2 | NMF-3 | NMF-4 | NMF-5 | NMF-6 | NMF-7 | NMF-8 | NMF-9 | NMF-10 |
|---|---|---|---|---|---|---|---|---|---|---|
| 1 | IL32 | ARHGAP15 | CD8B | CD2 | CD52 | PTPRC | CORO1A | LTB | HLA-DQA1 | IL2RA |
| 2 | CD3D | ITK | CD8A | ISG15 | CD3D | CD96 | ARHGDIB | CD52 | HLA-DPA1 | TNFRSF4 |
| 3 | CD52 | CD69 | GZMA | IL2RA | LTB | THEMIS | TYMS | IL32 | HLA-DPB1 | IL32 |
| 4 | S100A4 | SYTL3 | CD3D | CD3D | IL7R | LCP1 | RRM2 | S100A4 | HLA-DRB1 | TNFRSF18 |
| 5 | CD3E | BCL11B | IL32 | CYTIP | GIMAP7 | STK17B | IL32 | EMP3 | HLA-DRA | FOXP3 |
| 6 | CD2 | SRGN | NKG7 | COTL1 | CD3E | DOCK10 | HMGB2 | CORO1A | HLA-DRB5 | CTLA4 |
| 7 | FXYD5 | CHST11 | CD3E | IFI6 | LAPTM5 | MBNL1 | RAC2 | FXYD5 | SRGN | CD3D |
| 8 | CRIP1 | CNOT6L | CD52 | CORO1B | ARHGDIB | CLEC2D | MKI67 | IL2RG | CD74 | SRGN |
| 9 | HCST | IL7R | HCST | SRGN | HLA-B | CD2 | HLA-A | SH3BGRL3 | EBI3 | CD2 |
| 10 | IL2RG | CXCR4 | CORO1A | HLA-B | LCK | CD3G | LCK | ARHGDIB | CCL17 | TNFRSF1B |

**b**

Attribution-based NMF Top Genes per Factor (Highly Variable Genes)

| | NMF-1 | NMF-2 | NMF-3 | NMF-4 | NMF-5 | NMF-6 | NMF-7 | NMF-8 | NMF-9 | NMF-10 |
|---|---|---|---|---|---|---|---|---|---|---|
| 1 | ITK | CD8B | CD52 | GZMA | CD40LG | IL7R | ISG15 | HLA-DQA1 | IL2RA | PTPRC |
| 2 | ARHGAP15 | CD8A | SH3BGRL3 | NKG7 | HMGB2 | KLRB1 | CYTIP | HLA-DPA1 | TNFRSF4 | IKZF1 |
| 3 | SYTL3 | CD7 | CRIP1 | GZMB | SRGN | CD40LG | IL2RA | HLA-DPB1 | TNFRSF18 | STK17B |
| 4 | CD69 | CD96 | HLA-B | GZMH | CRIP1 | SRGN | CCR7 | HLA-DRB1 | FOXP3 | MBNL1 |
| 5 | CHST11 | GZMA | SELL | CST7 | RRM2 | HLA-B | HLA-B | HLA-DRA | SRGN | DOCK10 |
| 6 | CNOT6L | CD27 | ARHGAP15 | IFNG | MKI67 | CD96 | IFI6 | HLA-DRB5 | CTLA4 | BCL11B |
| 7 | BCL11B | NKG7 | CD27 | LAG3 | HMGN2 | CD69 | SRGN | SRGN | TNFRSF1B | APBB1IP |
| 8 | THEMIS | CD52 | IKZF1 | CD52 | CD96 | CXCR4 | TNFAIP8 | CCL17 | BATF | CD247 |
| 9 | FYN | THEMIS | CD44 | CD96 | SH3BGRL3 | ZFP36L2 | RPS17 | EBI3 | ICOS | STK4 |
| 10 | STK17B | CTSW | STK17B | CD8A | ALOX5AP | ANXA1 | MX1 | CCL22 | CD247 | ETS1 |

**c**

Expression-based NMF Top Genes per Factor (All Genes)

| | NMF-1 | NMF-2 | NMF-3 | NMF-4 | NMF-5 | NMF-6 | NMF-7 | NMF-8 | NMF-9 | NMF-10 |
|---|---|---|---|---|---|---|---|---|---|---|
| 1 | TMSB10 | MALAT1 | GZMA | MALAT1 | IL32 | ISG15 | GAPDH | TUBA1B | HSP90AB1 | IL6ST |
| 2 | ACTB | MT-CO2 | NKG7 | RPS27 | LTB | IFI6 | RPS2 | STMN1 | NPM1 | MALAT1 |
| 3 | RPS26 | MT-CO1 | GZMB | RPL13A | B2M | TMSB4X | RPS18 | UBE2C | HSPD1 | RPS27 |
| 4 | RPS27 | MT-ATP6 | GNLY | RPL41 | MT-ND4L | MX1 | EEF1A1 | HMGB2 | HSP90AA1 | B2M |
| 5 | TMSB4X | MT-ND2 | CD74 | RPL34 | ACTB | LGALS1 | ENO1 | TUBB | RPS2 | RPS29 |
| 6 | RPS28 | MT-ND3 | CST7 | EEF1A1 | EEF1A1 | S100A11 | RPLP1 | MKI67 | GAPDH | LINC-PINT |
| 7 | RPL28 | MT-ND4L | HLA-DRB1 | RPS18 | MT-CO2 | LY6E | DDIT4 | HMGN2 | PTMA | RBFOX2 |
| 8 | RPLP1 | MT-CYB | B2M | RPS29 | TMSB4X | ISG20 | LDHA | RRM2 | RPLP0 | STEAP1B |
| 9 | RPL37 | MT-ND1 | HLA-DRB5 | RPL13 | MT-CO1 | B2M | BNIP3 | TOP2A | C1QBP | TMSB4X |
| 10 | RPS8 | PTPRC | ACTB | RPL21 | RPL10 | MYL6 | EIF1 | CKS2 | HSPE1 | RPL34 |

**d**

Expression-based NMF Top Genes per Factor (Highly Variable Genes)

| | NMF-1 | NMF-2 | NMF-3 | NMF-4 | NMF-5 | NMF-6 | NMF-7 | NMF-8 | NMF-9 | NMF-10 |
|---|---|---|---|---|---|---|---|---|---|---|
| 1 | STEAP1B | MT-ND2 | HLA-DRB1 | TUBA1B | RPL31 | GZMA | ISG15 | LINC-PINT | IL2RA | CD52 |
| 2 | RBFOX2 | MT-ND3 | HLA-DPA1 | STMN1 | RPS20 | NKG7 | IFI6 | RBPJ | SAT1 | SH3BGRL3 |
| 3 | RPL17-C18orf32 | MT-ATP8 | HLA-DRA | TUBB | RPS17 | GZMB | MX1 | ANKRD28 | CYTIP | MT-ATP8 |
| 4 | IL6ST | SMCHD1 | HLA-DRB5 | HMGB2 | MT-ND2 | GNLY | HLA-B | ATXN1 | DDIT4 | HLA-B |
| 5 | ARMC9 | MBNL1 | HSP90AA1 | UBE2C | RPL36A | CD8A | RPS17 | FKBP5 | NEAT1 | FTH1 |
| 6 | RPL31 | LINC-PINT | MT-ATP8 | HMGN2 | HLA-B | SH3BGRL3 | ISG20 | RNF19A | SRGN | IFITM2 |
| 7 | RPS17 | FOXP1 | HLA-DPB1 | RRM2 | MT-ND3 | CD52 | RPS20 | AKAP13 | DUSP4 | MT-ND3 |
| 8 | RPL36A | PTPRC | FTH1 | MKI67 | FTH1 | CTSW | MT-ND2 | PARP8 | BTG1 | CRIP1 |
| 9 | RPS20 | CNOT6L | GSTP1 | CKS2 | ZFAS1 | CST7 | ZBED2 | SYTL3 | NFKBIA | IER2 |
| 10 | TXNRD1 | ARHGAP15 | HLA-DQA1 | TUBB4B | BTG1 | ALOX5AP | HSP90AA1 | MBNL1 | NAMPT | SARAF |

**e**

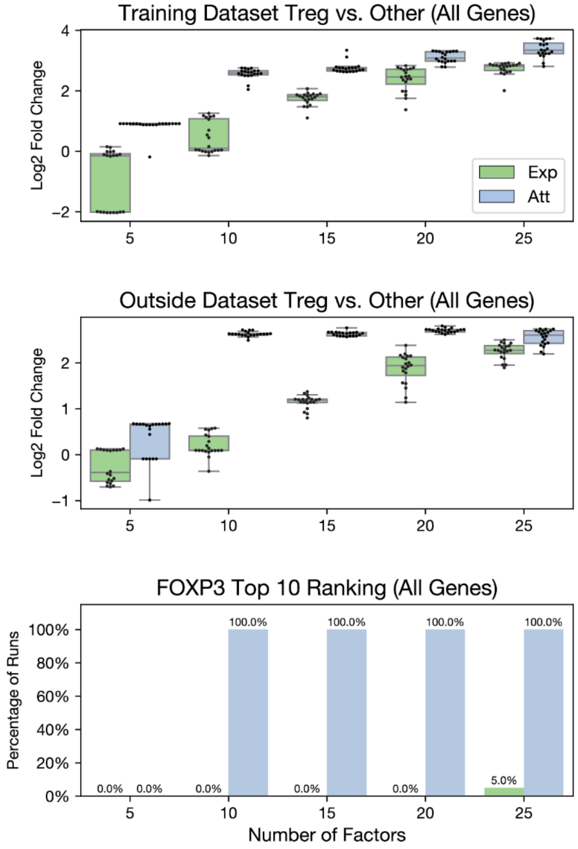


**f**

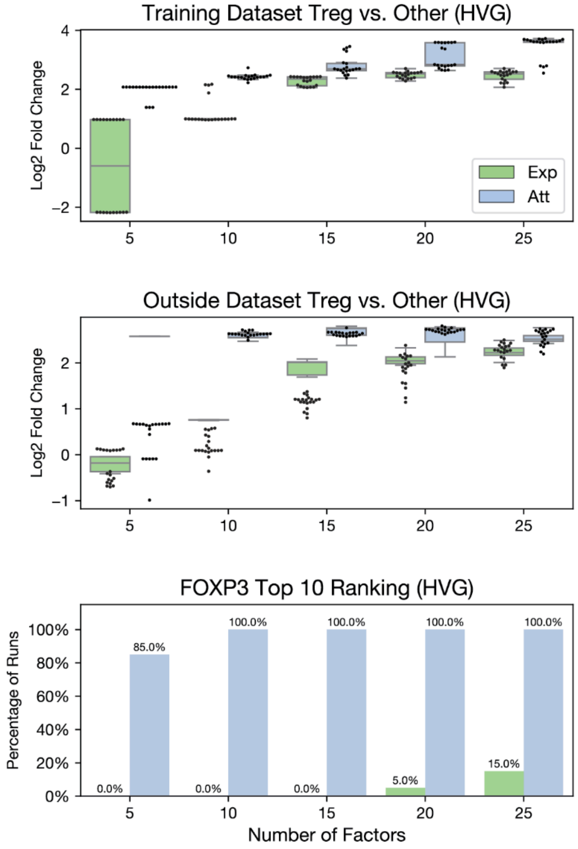

**<u>Extended Data Fig. 4</u>: NMF gene programs and generalization results**

Top 10 weighted genes produced by running NMF on T cells from across from three studies using 15 factors and random state of 114. The x-axis indicates the factor and the y-axis shows the ranking. **A.** Attribution-based NMF with all 14056 genes. **b.** Expression-based NMF with all 14056 genes. **c.** Attribution-based NMF with 2591 highly variable genes. **d.** Expression-based NMF with 2591 highly variable genes. **e.** shows robustness analysis using all 14056 genes , while **f.** shows the same results when using 2591 highly variable genes. In each case, NMF was applied and a single factor that best distinguishes Tregs from other T cells was chosen for downstream analysis (see Methods). The top box shows the log2 fold change between the Tregs and other T cells on the y-axis. The x-axis indicates the number of factors. The boxplots show data quartiles and each dot represents an individual run with a set random seed. The middle plot compares Tregs and other CD4+ T cells from 16 tissues not included in the original training dataset. The NMF models were applied to this data and the average usage was calculated for Tregs and other T cells for each tissue. Each dot represents the average usage for a cell type (Treg or CD4+ T cell) in a given tissue. The y-axis shows the log2 fold change between the tissue-level Treg values and the tissue-level CD4+ T cell values. The bottom plot shows the percentage of runs for that given set of parameters where FOXP3 appears in the top 10 genes.

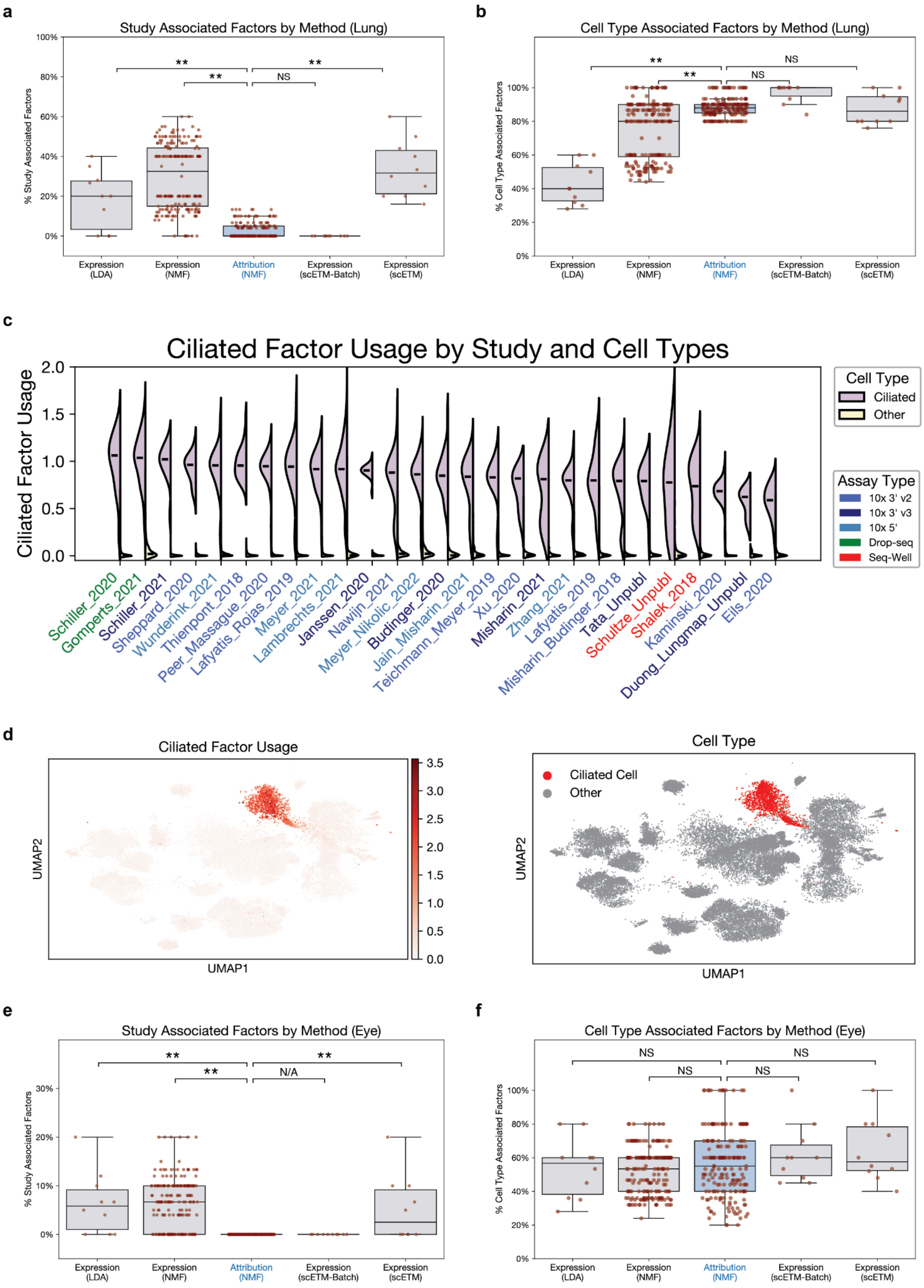
a
Study Associated Factors by Method (Lung)
% Study Associated Factors
100%
80%
60%
40%
20%
0%
**
**
NS
**
Expression (LDA)
Expression (NMF)
Attribution (NMF)
Expression (scETM-Batch)
Expression (scETM)
b
Cell Type Associated Factors by Method (Lung)
% Cell Type Associated Factors
100%
80%
60%
40%
20%
0%
**
NS
**
NS
Expression (LDA)
Expression (NMF)
Attribution (NMF)
Expression (scETM-Batch)
Expression (scETM)
c
Ciliated Factor Usage by Study and Cell Types
Ciliated Factor Usage
2.0
1.5
1.0
0.5
0.0
Cell Type
Ciliated
Other
Assay Type
10x 3' v2
10x 3' v3
10x 5'
Drop-seq
Seq-Well
Schiller_2020
Gomperts_2021
Schiller_2021
Sheppard_2020
Wunderink_2021
Thienpont_2018
Peer_Massague_2020
Lafyatis_Rojas_2019
Meyer_2021
Lambrechts_2021
Janssen_2020
Nawijn_2021
Meyer_Nikolic_2022
Budinger_2020
Jain_Misharin_2021
Teichmann_Meyer_2019
Xu_2020
Misharin_2021
Zhang_2021
Lafyatis_2019
Misharin_Budinger_2018
Tata_Unpubl
Schultze_Unpubl
Shalek_2018
Kaminski_2020
Duong_Lungmap_Unpubl
Eils_2020
d
Ciliated Factor Usage
UMAP2
UMAP1
3.5
3.0
2.5
2.0
1.5
1.0
0.5
0.0
Cell Type
Ciliated Cell
Other
UMAP2
UMAP1
e
Study Associated Factors by Method (Eye)
% Study Associated Factors
30%
20%
10%
0%
**
**
N/A
**
Expression (LDA)
Expression (NMF)
Attribution (NMF)
Expression (scETM-Batch)
Expression (scETM)
f
Cell Type Associated Factors by Method (Eye)
% Cell Type Associated Factors
100%
80%
60%
40%
20%
0%
NS
NS
NS
NS
Expression (LDA)
Expression (NMF)
Attribution (NMF)
Expression (scETM-Batch)
Expression (scETM)

**<u>Extended Data Fig. 5</u>: Unsupervised learning on lung and eye scRNA-seq data**
Each dot in the top panels represents a parameter set used for NMF, LDA, scETM, or supervised scETM using batch labels (scETM-batch). The x-axis indicates the matrix values (attributions or expression) and the method (NMF, LDA, scETM, scETM-batch). The y-axis indicates the percentage of factors or topics in that run that are correlated with either cell type or study (adjusted $r^2 > 0.25$). Lung data is from the Human Lung Cell Atlas [4], whereas ocular corneal data is downloaded from the CZI database [5]. **a.** lung data correlated with study, **b.** lung data correlated with cell type. **c.** Violin plot showing NMF factor 15 usage in ciliated cells (purple) and other cells (yellow) for each study. The values The sequencing platform used by each study is shown in the legend on the bottom right. Ciliated cell counts: 60, 115, 35, 37, 63, 36, 38, 34, 55, 231, 25, 100, 315, 30, 70, 19, 11, 16, 80, 25, 100, 28, 16, 119, 394, 10, 58. Other cell counts: 1560, 531, 1008, 1149, 839, 1221, 951, 762, 1273, 2020, 517, 1251, 1900, 566, 529, 555, 292, 346, 775, 601, 1911, 449, 219, 1318, 8640, 176, 916. **d.** UMAP showing NMF factor 15 usage (left) and ciliated cell positions (right) using the original UMAP coordinates from the human lung cell atlas [4]. **e.** eye data correlated with study, **f.** eye data correlated with cell type. Each box indicates the data quartiles. ** indicates adj-p < 0.01 (Mann Whitney). “NS” indicates not significant. “N/A” indicates not applicable, indicating the statistical test could not be calculated as both distributions were all 0s. For each plot, expression and attribution NMF values have 200 parameter set dots and topic modeling with LDA or scETM have 10.

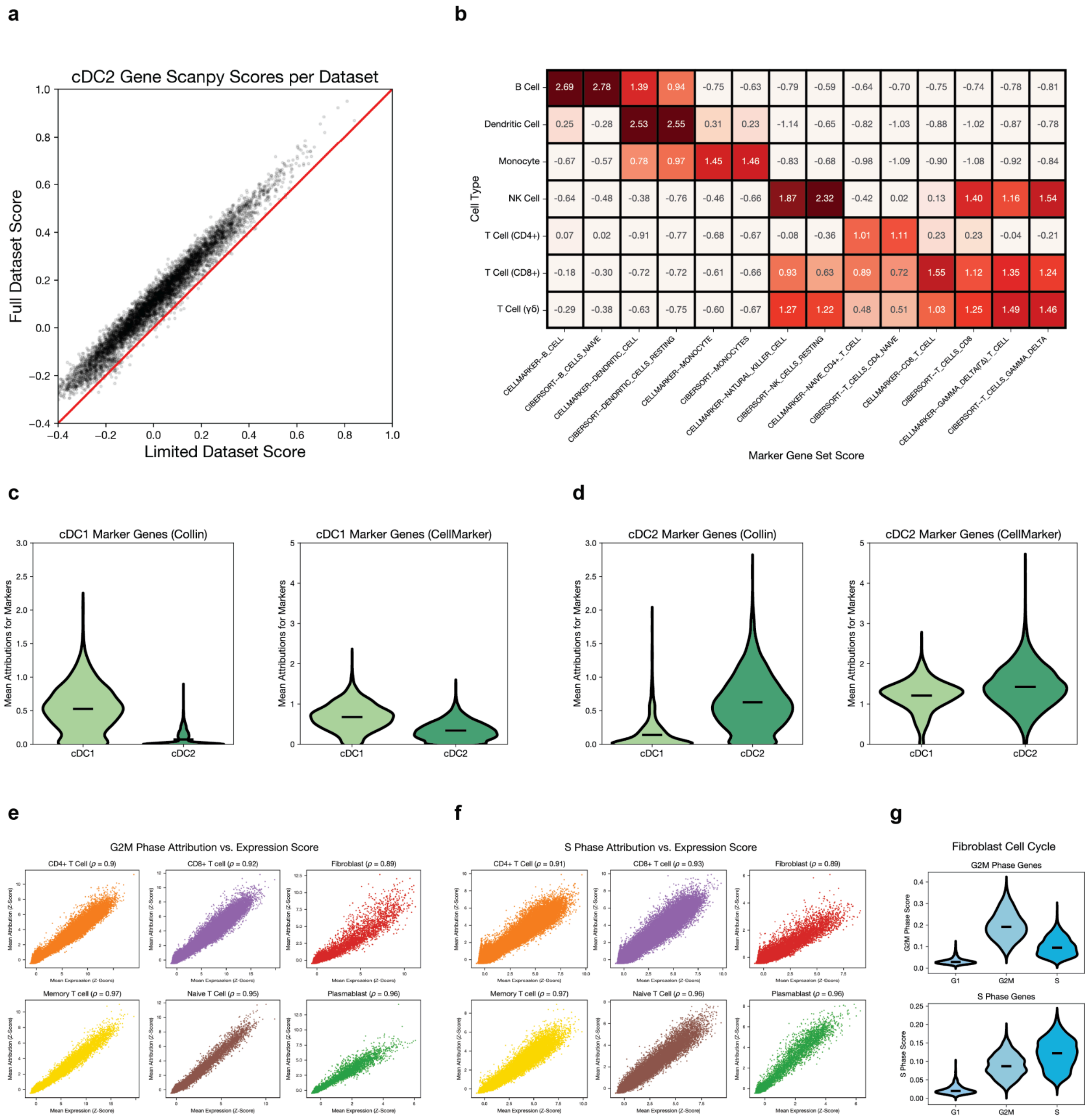
a
b
c
d
e
f
g
cDC2 Gene Scanpy Scores per Dataset
Full Dataset Score
Limited Dataset Score
Cell Type
B Cell
Dendritic Cell
Monocyte
NK Cell
T Cell (CD4+)
T Cell (CD8+)
T Cell (γδ)
CELLMARKER--B_CELL
CIBERSORT--B_CELLS_NAIVE
CELLMARKER--DENDRITIC_CELL
CIBERSORT--DENDRITIC_CELLS_RESTING
CELLMARKER--MONOCYTE
CIBERSORT--MONOCYTES
CELLMARKER--NATURAL_KILLER_CELL
CIBERSORT--NK_CELLS_RESTING
CELLMARKER--NAIVE_CD4+_T_CELL
CIBERSORT--T_CELLS_CD4_NAIVE
CELLMARKER--CD8_T_CELL
CIBERSORT--T_CELLS_CD8
CELLMARKER--GAMMA_DELTA(Δ)_T_CELL
CIBERSORT--T_CELLS_GAMMA_DELTA
Marker Gene Set Score
2.69 2.78 1.39 0.94 -0.75 -0.63 -0.79 -0.59 -0.64 -0.70 -0.75 -0.74 -0.78 -0.81
0.25 -0.28 2.53 2.55 0.31 0.23 -1.14 -0.65 -0.82 -1.03 -0.88 -1.02 -0.87 -0.78
-0.67 -0.57 0.78 0.97 1.45 1.46 -0.83 -0.68 -0.98 -1.09 -0.90 -1.08 -0.92 -0.84
-0.64 -0.48 -0.38 -0.76 -0.46 -0.66 1.87 2.32 -0.42 0.02 0.13 1.40 1.16 1.54
0.07 0.02 -0.91 -0.77 -0.68 -0.67 -0.08 -0.36 1.01 1.11 0.23 0.23 -0.04 -0.21
-0.18 -0.30 -0.72 -0.72 -0.61 -0.66 0.93 0.63 0.89 0.72 1.55 1.12 1.35 1.24
-0.29 -0.38 -0.63 -0.75 -0.60 -0.67 1.27 1.22 0.48 0.51 1.03 1.25 1.49 1.46
cDC1 Marker Genes (Collin)
cDC1 Marker Genes (CellMarker)
cDC2 Marker Genes (Collin)
cDC2 Marker Genes (CellMarker)
Mean Attributions for Markers
cDC1
cDC2
G2M Phase Attribution vs. Expression Score
S Phase Attribution vs. Expression Score
CD4+ T Cell (ρ = 0.9)
CD8+ T cell (ρ = 0.92)
Fibroblast (ρ = 0.89)
Memory T cell (ρ = 0.97)
Naive T Cell (ρ = 0.95)
Plasmablast (ρ = 0.96)
CD4+ T Cell (ρ = 0.91)
CD8+ T cell (ρ = 0.93)
Fibroblast (ρ = 0.89)
Memory T cell (ρ = 0.97)
Naive T Cell (ρ = 0.96)
Plasmablast (ρ = 0.96)
Mean Attribution (Z-Score)
Mean Expression (Z-Score)
Fibroblast Cell Cycle
G2M Phase Genes
G2M Phase Score
S Phase Genes
S Phase Score
G1
G2M
S

**<u>Extended Data Fig. 6</u>: Gene Set Scoring**

**a.** Each dot represents a single conventional dendritic cell (type 2) from Adams *et al* [3]. The y-axis indicates the Scanpy score using cDC2 marker genes when calculated on the full dataset. The x-axis shows the Scanpy score calculated using the same cDC2 marker genes but on a limited dataset only considering conventional dendritic cells and mast cells. **b.** Average attributions were calculated for seven categories of immune cells from Van Der Wijst et al. [6] and z-scored across cells. Gene sets were used from CIBERSORT [7] and CellMarker 2.0 [8]. The y-axis shows the cell type. The x-axis indicates the gene set source and specific gene set. The box shows the average z-score for all cells among the population indicated by the y-axis. **c-d.** Average attributions for marker genes for conventional dendritic cells, type 1 (cDC1, n=1122) **c.** and type 2 (cDC2, n=6760) **d.** X-axis indicates the cell population. Y-axis indicates the score for the marker genes indicated in the title. **e.** G2M phase marker scores from [9] across 6 cell types most present in the top 10000 scoring cells. The y-axis indicates average attributions for G2M marker genes and x-axis shows average log-normalized expression. **f.** Same results for S phase. **g.** Mean attributions for G2M markers (top) and S markers (bottom) for fibroblasts, whose cell cycle phases (x-axis) were inferred from Riba et al. [1] (2281 G1, 1567 S, 1519 G2M).

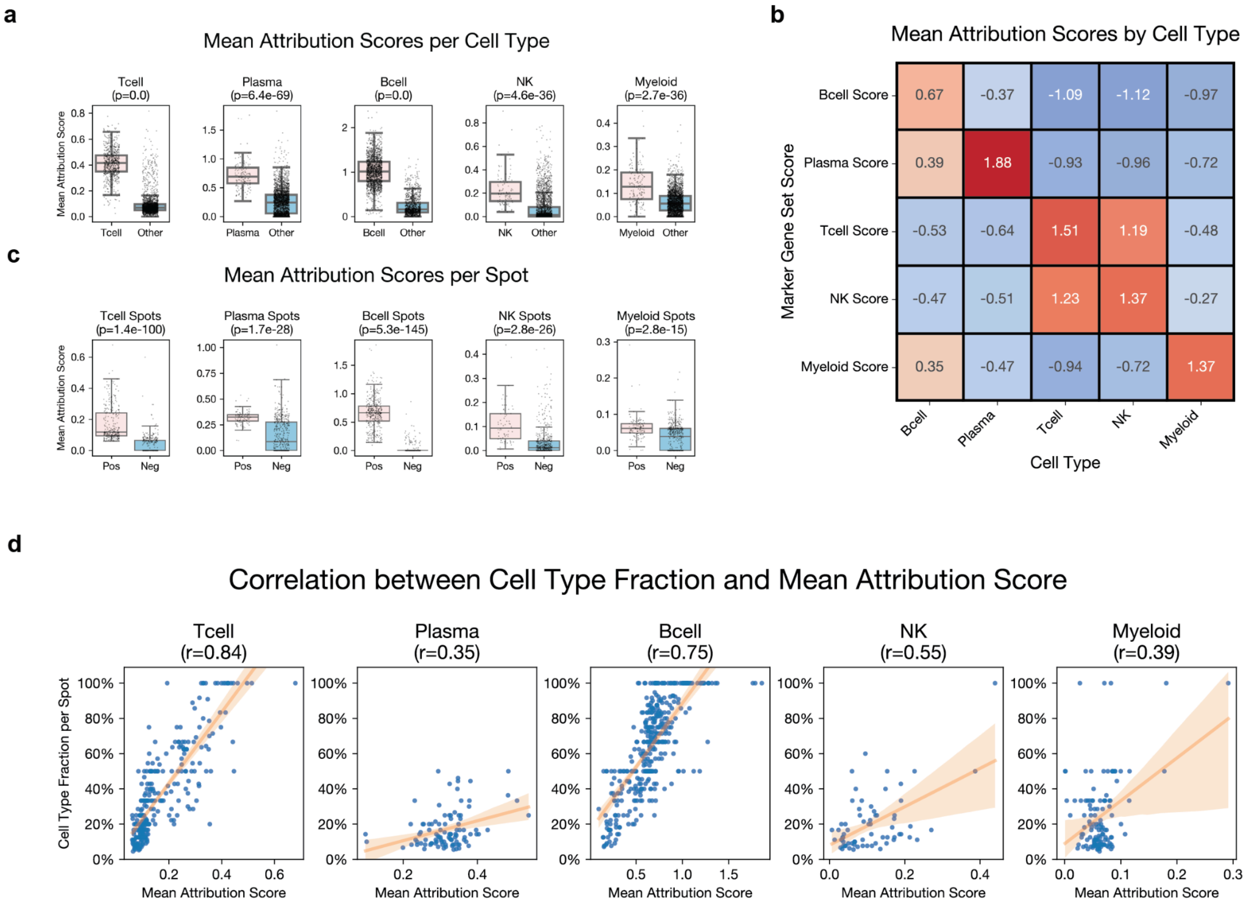


**<u>Extended Data Fig. 7</u>: Attribution calculation on spatial transcriptomics**

**a.** Mean attribution scores for cell type markers are compared across individual cells from Russell *et al.* [10] (n=3,119). The box plots show data quartiles. **b.** Heatmap shows the mean standardized gene attribution scores for a given marker gene set (y-axis) in a given cell population (x-axis). **c.** Mean attribution scores for cell type markers are compared across simulated spots (n=625) (**Methods**). **d.** The correlation between the mean attribution score for a cell type (x-axis) and its composition within each spot (y-axis) is shown. Only spots containing at least one example of the cell type of interest are displayed. Orange lines show the line of best fit, drawn using the regplot function from *seaborn* [11].

| Cell Type | Cell Type Info | | Mean Logistic Regression F-Score | | | | | | Otsu Thresholding F-Score | | | | | |
|---|---|---|---|---|---|---|---|---|---|---|---|---|---|---|
| | Num Cells | Pct Total | ANS | Jasmine | Mean Atttribution | Mean Expression | Scanpy | UCell | ANS | Jasmine | Mean Atttribution | Mean Expression | Scanpy | UCell |
| **ATI Epithelial Cell** | 1032 | 0.42% | 0.7368 | 0.5772 | **0.9025** | 0.5197 | 0.6711 | 0.5641 | 0.9194 | 0.0113 | **0.9582** | 0.7589 | 0.8327 | 0.8167 |
| **ATII Epithelial Cell** | 3881 | 1.57% | 0.8755 | 0.2636 | **0.9163** | 0.6318 | 0.7217 | 0.5371 | 0.8544 | 0.1801 | **0.8838** | 0.7638 | 0.7829 | 0.7480 |
| **B Cell** | 7798 | 3.16% | 0.9603 | 0.9102 | **0.9815** | 0.9282 | 0.9518 | 0.9272 | 0.9731 | 0.9581 | **0.9850** | 0.9440 | 0.9630 | 0.9438 |
| **Basal Cell** | 1670 | 0.68% | 0.0989 | 0.0109 | **0.2542** | 0.0229 | 0.0225 | 0.0182 | 0.1824 | 0.0106 | **0.2742** | 0.0121 | 0.0120 | 0.0116 |
| **Ciliated Cell** | 9361 | 3.80% | 0.9901 | 0.9904 | **0.9928** | 0.9824 | 0.9887 | 0.9892 | 0.9834 | 0.9886 | **0.9917** | 0.9597 | 0.9811 | 0.9808 |
| **Classical Monocyte** | 15391 | 6.24% | **0.5317** | 0.4574 | 0.5250 | 0.2164 | 0.3147 | 0.2415 | **0.6244** | 0.1396 | 0.1398 | 0.1494 | 0.1463 | 0.1477 |
| **Club Cell** | 2492 | 1.01% | 0.0470 | 0.0284 | **0.1390** | 0.0158 | 0.0157 | 0.0155 | 0.0828 | 0.0133 | **0.2907** | 0.0140 | 0.0141 | 0.0137 |
| **Conventional Dendritic Cell Type 1** | 1122 | 0.46% | 0.0366 | 0.0187 | **0.1159** | 0.0231 | 0.0257 | 0.0201 | 0.0393 | 0.0097 | **0.1408** | 0.0136 | 0.0136 | 0.0142 |
| **Fibroblast** | 2605 | 1.06% | **0.3188** | 0.0694 | 0.2186 | 0.2104 | 0.2712 | 0.1705 | **0.3952** | 0.0848 | 0.3593 | 0.3363 | 0.3090 | 0.2894 |
| **Goblet Cell** | 1262 | 0.51% | 0.4668 | 0.1509 | **0.5888** | 0.1987 | 0.3352 | 0.2458 | 0.6909 | 0.0130 | 0.6507 | 0.5695 | 0.6619 | **0.7047** |
| **Macrophage** | 178364 | 72.37% | 0.9314 | 0.9203 | **0.9370** | 0.9062 | 0.9124 | 0.9051 | 0.9047 | 0.9199 | **0.9374** | 0.8946 | 0.9060 | 0.8810 |
| **Mesothelial Cell** | 493 | 0.20% | **0.0153** | 0.0032 | 0.0079 | 0.0047 | 0.0047 | 0.0047 | 0.0047 | 0.0033 | **0.0048** | 0.0046 | 0.0036 | 0.0046 |
| **Myofibroblast** | 3477 | 1.41% | 0.5528 | 0.1388 | **0.6307** | 0.5149 | 0.5527 | 0.4758 | 0.6734 | 0.0295 | 0.6727 | 0.6543 | **0.6739** | 0.6487 |
| **Natural Killer Cell** | 11212 | 4.55% | 0.9462 | 0.9222 | **0.9668** | 0.9358 | 0.9392 | 0.9284 | 0.9552 | 0.9497 | **0.9671** | 0.9528 | 0.9511 | 0.9445 |
| **Non-classical Monocyte** | 5616 | 2.28% | **0.4240** | 0.3166 | 0.3691 | 0.1858 | 0.2781 | 0.2046 | **0.4418** | 0.0536 | 0.0538 | 0.0586 | 0.0564 | 0.0580 |
| **Plasmacytoid Dendritic Cell** | 691 | 0.28% | **0.9379** | 0.5915 | 0.8586 | 0.9258 | 0.9373 | 0.9066 | **0.9556** | 0.0137 | **0.9574** | 0.9510 | 0.9525 | 0.9449 |

**Extended Data Fig. 8: F-Scores for supervised and unsupervised classification tasks**

Left column shows the cell type. The next section indicates the number of cells and percentage of the dataset. The middle region shows the mean F-score using 25 train/test splits for supervised logistic regression predicting the cell population from its marker gene scores. The yellow bolded boxes indicate the top scoring method for a given cell type.The right panel shows the F-score for unsupervised classification using otsu thresholding. The gray bolded boxes indicate the top scoring method for a given cell type.

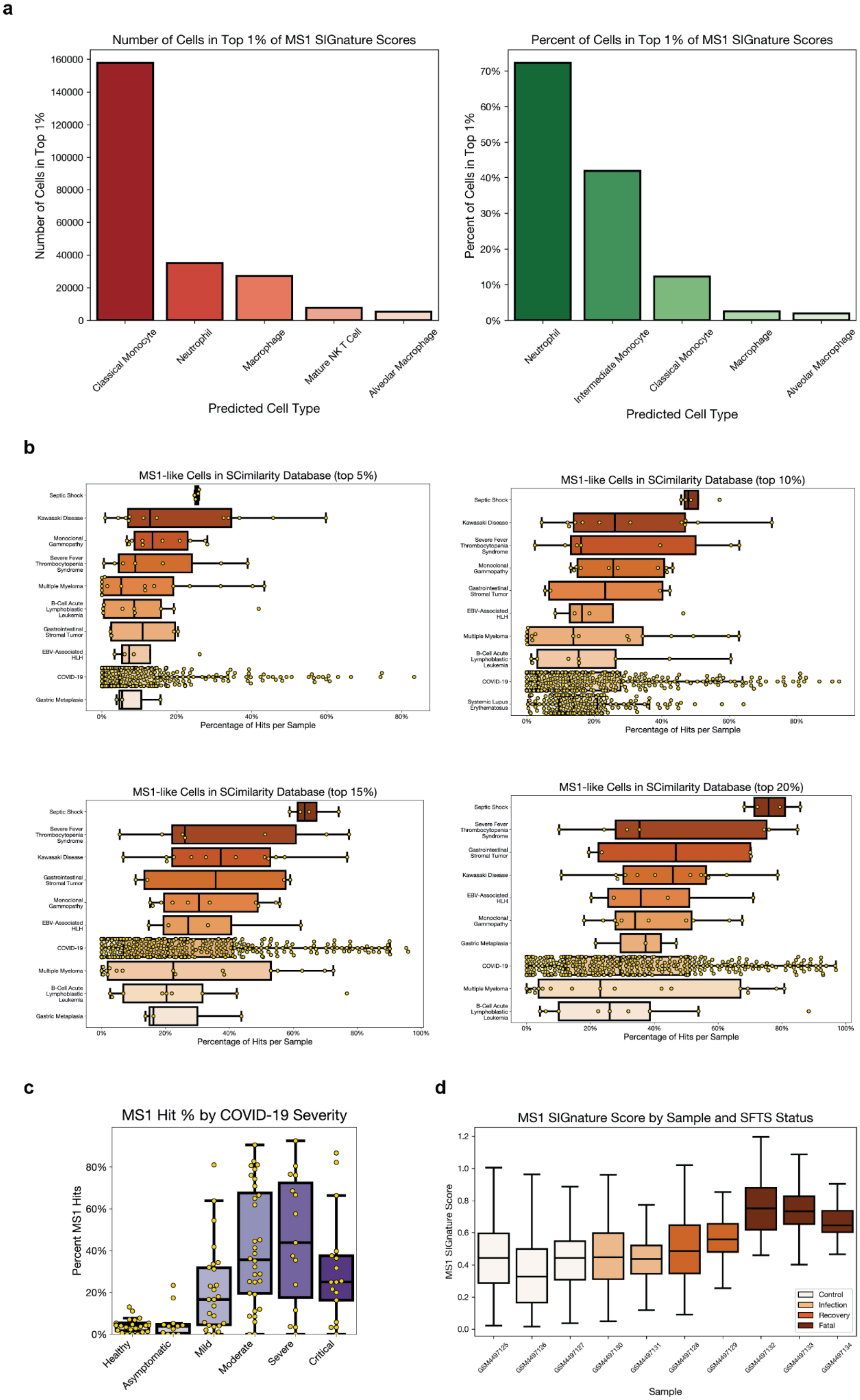
a
Number of Cells in Top 1% of MS1 SIGnature Scores
Number of Cells in Top 1%
160000
140000
120000
100000
80000
60000
40000
20000
0
Classical Monocyte
Neutrophil
Macrophage
Mature NK T Cell
Alveolar Macrophage
Predicted Cell Type
Percent of Cells in Top 1% of MS1 SIGnature Scores
Percent of Cells in Top 1%
70%
60%
50%
40%
30%
20%
10%
0%
Neutrophil
Intermediate Monocyte
Classical Monocyte
Macrophage
Alveolar Macrophage
Predicted Cell Type
b
MS1-like Cells in SCimilarity Database (top 5%)
Septic Shock
Kawasaki Disease
Monoclonal Gammopathy
Severe Fever Thrombocytopenia Syndrome
Multiple Myeloma
B-Cell Acute Lymphoblastic Leukemia
Gastrointestinal Stromal Tumor
EBV-Associated HLH
COVID-19
Gastric Metaplasia
Percentage of Hits per Sample
MS1-like Cells in SCimilarity Database (top 10%)
Septic Shock
Kawasaki Disease
Severe Fever Thrombocytopenia Syndrome
Monoclonal Gammopathy
Gastrointestinal Stromal Tumor
EBV-Associated HLH
Multiple Myeloma
B-Cell Acute Lymphoblastic Leukemia
COVID-19
Systemic Lupus Erythematosus
Percentage of Hits per Sample
MS1-like Cells in SCimilarity Database (top 15%)
Septic Shock
Severe Fever Thrombocytopenia Syndrome
Kawasaki Disease
Gastrointestinal Stromal Tumor
Monoclonal Gammopathy
EBV-Associated HLH
COVID-19
Multiple Myeloma
B-Cell Acute Lymphoblastic Leukemia
Gastric Metaplasia
Percentage of Hits per Sample
MS1-like Cells in SCimilarity Database (top 20%)
Septic Shock
Severe Fever Thrombocytopenia Syndrome
Gastrointestinal Stromal Tumor
Kawasaki Disease
EBV-Associated HLH
Monoclonal Gammopathy
Gastric Metaplasia
COVID-19
Multiple Myeloma
B-Cell Acute Lymphoblastic Leukemia
Percentage of Hits per Sample
c
MS1 Hit % by COVID-19 Severity
Percent MS1 Hits
80%
60%
40%
20%
0%
Healthy
Asymptomatic
Mild
Moderate
Severe
Critical
d
MS1 SIGnature Score by Sample and SFTS Status
MS1 SIGnature Score
1.2
1.0
0.8
0.6
0.4
0.2
0.0
GSM4497125
GSM4497126
GSM4497127
GSM4497130
GSM4497131
GSM4497128
GSM4497129
GSM4497132
GSM4497133
GSM4497134
Sample
Control
Infection
Recovery
Fatal

**<u>Extended Data Fig. 9:</u> Querying Attributions for MS1 gene signature**

**a.** Cell types enriched in top 1% of MS1 scores. Left barplot shows the top 5 cell types by number. Right barplot shows the top 5 cell types by proportion. **b.** Top MS1-associated diseases by hit threshold. The y-axis indicates disease and the x-axis shows the percentage of monocytes/macrophages per sample that are above hit threshold. Boxplots show data quartiles and yellow dots indicate individual samples. Each subplot displays a different threshold, indicated in the title. **c.** Percentage of MS1-like cells by COVID severity from Stephenson *et al.* [12] . X-axis indicates COVID-19 severity. Y-axis shows the percentage of MS1-like monocytes or macrophages in each sample. Boxplots show data quartiles. **d.** MS1 scores in SFTS cells by severity. The y-axis indicates mean MS1 SIGnature score for monocytes in SFTS dataset [13]. X-axis indicates the sample name. Boxplots are colored by SFTS status, shown in legend in top left and show the data quartiles for monocytes in that sample. Cells outside the 1.5 IQR range are not shown. The boxes represent the following cell counts: 1140, 797, 1196, 663, 938, 326, 266, 60, 662, 98.

a

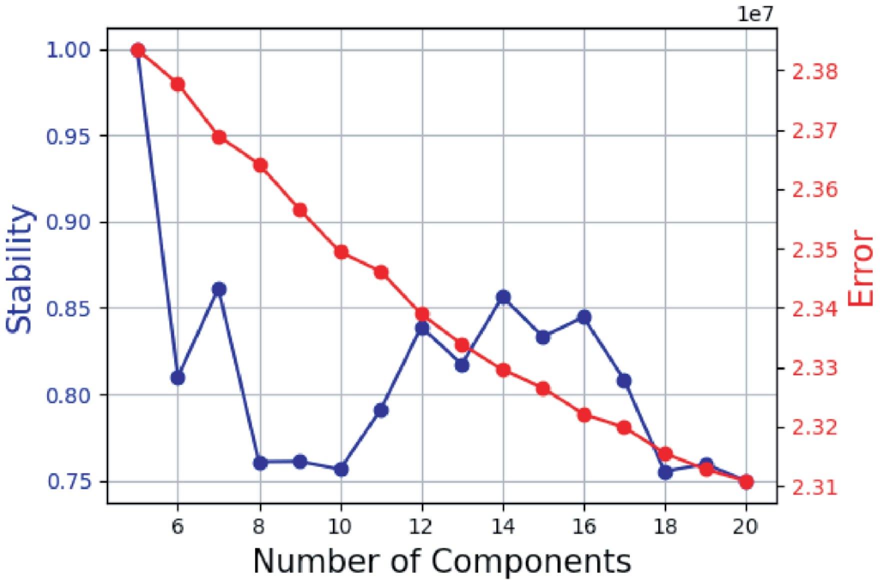


b

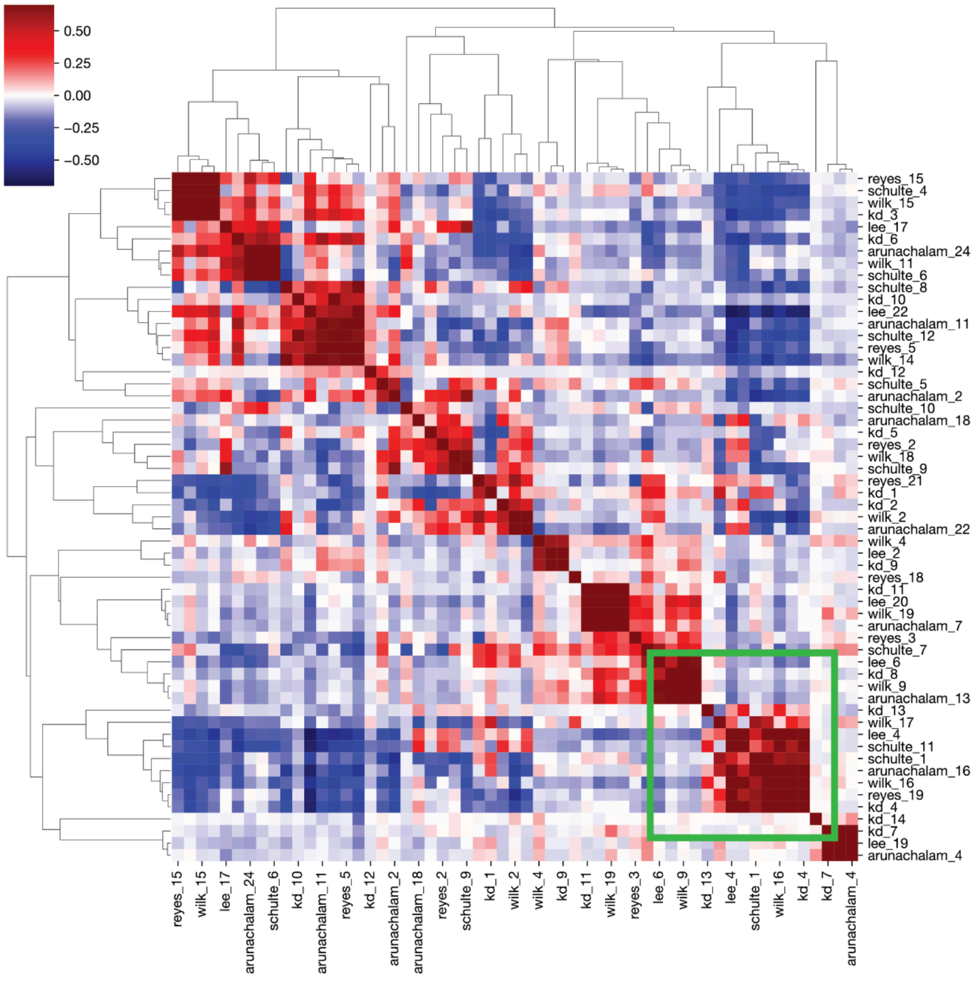

**<u>Extended Data Figure 10</u>: Kawasaki disease analysis**

**a.** Consensus NMF factor parameters. Consensus NMF was run for components 5 through 20. The left y-axis in blue shows component stability over iterations. The right y-axis in red shows the reconstruction error from the NMF. 14 components were chosen as the peak stability after the initial decline as a way to optimize high stability and low error. **b.** Clustermap between standardized gene loadings from consensus NMF. Each row represents an individual factor from a study. The "Reyes", "Schulte", "Wilk", "Lee", and "Arunachalam" all indicate studies processed and analyzed in Reyes *et al.* [14]. The "KD" label indicates factors from this current study. The normalized gene loadings were compared using Pearson correlation and shown with a seaborn clustermap (**Methods**). The color bar at the top left shows the strength and direction of the correlation with deeper red being positive and deeper blue indicating negative. The green box highlights the area shown in Figure 4f.
